# Prediction-error neurons in circuits with multiple neuron types: Formation, refinement and functional implications

**DOI:** 10.1101/2021.08.24.457531

**Authors:** Loreen Hertäg, Claudia Clopath

## Abstract

Predictable sensory stimuli do not evoke significant responses in a subset of cortical excitatory neurons. Some of those neurons, however, change their activity upon mismatches between actual and predicted stimuli. Different variants of these prediction-error neurons exist and they differ in their responses to unexpected sensory stimuli. However, it is unclear how these variants can develop and co-exist in the same recurrent network, and how they are simultaneously shaped by the astonishing diversity of inhibitory interneurons. Here, we study these questions in a computational network model with three types of inhibitory interneurons. We find that balancing excitation and inhibition in multiple pathways gives rise to heterogeneous prediction-error circuits. Dependent on the network’s initial connectivity and distribution of actual and predicted sensory inputs, these circuits can form different variants of prediction-error neurons that are robust to network perturbations and generalize to stimuli not seen during learning. These variants can be learned simultaneously via homeostatic inhibitory plasticity with low baseline firing rates. Finally, we demonstrate that prediction-error neurons can support biased perception, we illustrate a number of functional implications, and we discuss testable predictions.

## Introduction

The theory of predictive processing posits that neural networks strive to predict sensory inputs and use prediction errors to constantly refine an inner model of the world (1–3). Neural hallmarks of prediction errors have been found widely. Dopaminergic neurons in the basal ganglia and striatum encode reward-prediction errors (4). Some neurons in layer 2/3 of rodent primary visual cortex (V1) (5, 6), or neurons in the telencephalic areas of adult zebrafish (7) are driven by mismatches between actual and predicted visual consequences of motor commands. Similarly, a subset of cortical neurons responds to auditory feedback perturbations during vocalization (8, 9), and some excitatory cells in mouse barrel cortex are sensitive to abrupt mismatches of tactile flow and the animal’s running speed (10). However, those neurons are embedded in complex circuits that exhibit a rich diversity of cell types interacting in many ways (11–15). It is mostly unresolved whether and how this diversity collaboratively shapes, processes, and refines prediction errors.

Mismatches may occur in two variants: sensory inputs can be overpredicted or underpredicted, depending on whether the prediction is larger or smaller than the sensory stimulus, respectively. While dopaminergic neurons signal mismatches bidirectionally (4), this may be impossible for neurons with very low spontaneous firing rates, as for instance cortical neurons in layer 2/3 of V1 (16, 17), because negative deviations would be bounded from below. Thus, it has been suggested that cortical prediction-error neurons come in two flavors (1, 3): negative prediction-error (nPE) neurons only increase their activity relative to baseline when a sensory stimulus is smaller than predicted, while positive prediction-error (pPE) neurons only increase activity when a sensory stimulus is larger than predicted.

Computing prediction errors, no matter whether negative or positive mismatches, requires inhibition (3). Despite being outnumbered by excitatory neurons, inhibitory interneurons shape cortical computations in many ways (11, 14, 18–22). This rich repertoire of interneuron function is accompanied by a great diversity in their morphology, physiology, connectivity patterns and synaptic properties (11, 14). In a computational model of layer 2/3 of rodent V1, it has been shown that the presence of nPE neurons imposes constraints on the interneuron network in the form of a balance of excitation and inhibition (23). However, it is not resolved whether the co-existence of nPE and pPE neurons imposes further requirements on the interneuron circuit they are embedded in. Moreover, the formation of mismatch neurons in layer 2/3 of V1 relies on normal visuomotor coupling during development (6). This suggests that PE neurons are experience-dependent, raising the question of how networks self-organize to give rise to them. While it has been shown that separate nPE and pPE circuits can be learned by means of homoestatic inhibitory plasticity (23), it is not resolved if and how this generalizes to networks with both nPE and pPE neurons.

To elucidate the circuit-level mechanisms that underlie the formation of both PE neuron types, we design a rate-based computational model with excitatory and three types of inhibitory neurons. We first show that in a simplified mean-field network, nPE and pPE neurons can co-exist when a balance of excitation and inhibition (E/I balance) is established. Moreover, the interneuron circuit must comprise two distinct sources of somatic inhibition. The dendritic inhibition must be driven by feedforward bottom-up signals. Once established, these PE neurons are robust to moderate network perturbations. We then simulate a heterogeneous network model and show that both nPE and pPE neurons can be learned simultaneously by inhibitory homeostatic plasticity when the network is exposed to predicted sensory inputs and the excitatory neurons exhibit low baseline firing rates. When synaptic plasticity establishes a balance of excitatory and inhibitory inputs, the PE neurons are robust and generalize to stimuli not seen during learning. Furthermore, we show that the formation of PE neurons depends on synaptic noise, the distribution of actual and predicted sensory inputs, and the initial connectivity between neurons. Finally, we connect a heterogeneous PE circuit with an attractor network and show that PE neurons can support biased perception, a phenomenon observed in tasks that involve the reproduction of a perceived variable (24–29). By means of the example of a contraction bias, we illustrate a number of functional implications for PE neurons. We show that they can act as an internal cue switching the network between attractors, may underpin generalization across distinct stimuli statistics, and can support faster learning.

## Results

Given that neural circuits contain an astonishing variety of neuron types and cell type-specific connections (11, 13, 14), we wondered under which constraints both negative and positive prediction-error (nPE/pPE) neurons can develop simultaneously in the same recurrent network. To address this question, we studied a rate-based network model with excitatory pyramidal cells (PCs), and inhibitory parvalbumin-expressing (PV), somatostatin-expressing (SOM), and vasoactive intestinal peptide-expressing (VIP) interneurons (Fig. 1A). The relative distribution of neuron types, their connection probabilities and strengths are motivated by electrophysiological studies (e.g., 12, 13, 30–35, see Methods for details). While all inhibitory neurons are modeled as point neurons (36), the excitatory neurons are simulated as two coupled point compartments, representing the soma and the dendrites, respectively.

**Figure 1.**
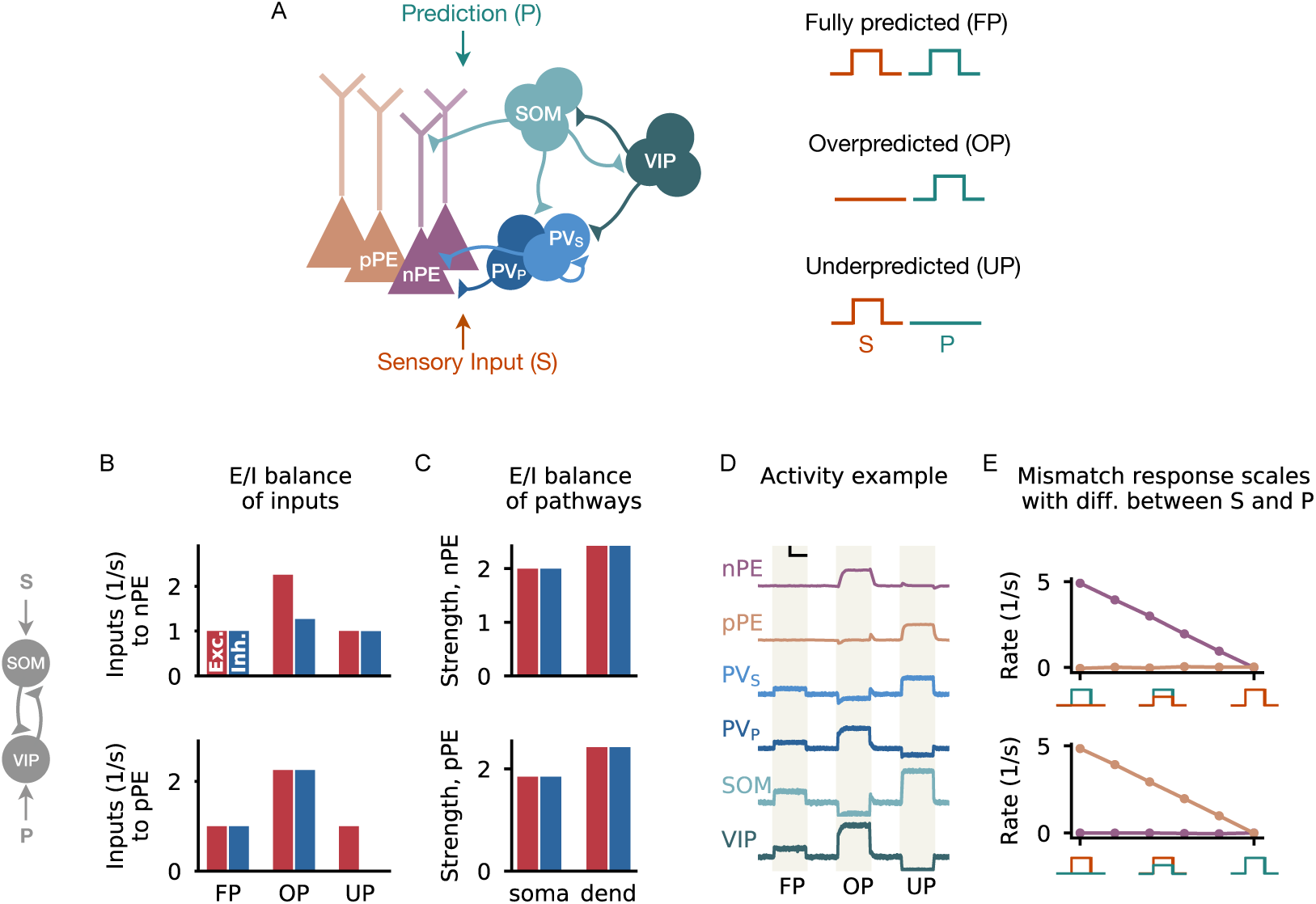
Multi-pathway E/I balance in nPE and pPE neurons. **(A)** Left: Network model with excitatory PCs and inhibitory PV, SOM and VIP neurons. Connections from PCs not shown for the sake of clarity. In a PE circuit, PCs either act as nPE or pPE neurons. Somatic compartment of PCs and half of the PV neurons (PV_S_) receive the actual sensory input, while the dendritic compartment of PCs and the remaining PV neurons (PV_P_) receive the predicted sensory input. SOM and VIP neurons receive either actual or predicted sensory input. Right: Sensory stimuli can either be fully predicted (FP), overpredicted (OP) or underpredicted (UP). **(B-E)** Mean-field network derived from (A) with the SOM neuron receiving the actual sensory input and the VIP neuron receiving the predicted sensory stimulus. **(B)** In a PE circuit, the excitatory (red) and inhibitory (blue) inputs are balanced for fully predicted sensory stimuli for both nPE and pPE neurons. This balance is preserved for underpredicted stimuli (nPE neurons, top) or overpredicted stimuli (pPE neurons, bottom). Stimulus strength is 1 *s*^−1^. **(C)** Both nPE (top) and pPE neurons (bottom) exhibit balanced pathways onto both soma and dendrites. **(D)** Example activity of all neuron types for fully predicted as well as over- and underpredicted stimuli for a network parameterized to establish an E/I balance in the pathways. Vertical black bar denotes 3 *s*^−1^, horizontal black bar denotes 500 ms. **(E)** Mismatch responses for nPE (top) and pPE (bottom) neurons scale with the difference between actual and predicted sensory inputs.

All neurons receive an excitatory background input to ensure reasonable baseline firing rates in the absence of any sensory stimulation (‘baseline phase’). In addition, we stimulated the neurons with time-varying inputs that represent actual and predicted sensory stimuli. It is known that excitatory neurons receive feedforward, sensory inputs at their basal dendrites and perisomatic region, and feedback projections from higher-order cortical areas at their apical dendrites (37, 38). These feedback projections are assumed to carry information about expectations, beliefs, or predictions (37, 39, 40), and mediate a broad range of functional roles (41, 42). While the distribution of feedforward and feedback inputs among the compartments of PCs is well-studied, the distribution among different types of cortical inhibitory interneurons is less certain and likely diverse (14, 38, 43, 44). To account for different input distributions and their effect on the formation of nPE and pPE neurons, the inputs onto inhibitory interneurons are varied in our simulations.

To identify nPE and pPE neurons, we modeled their responses to different combinations of actual and predicted sensory information. When the sensory input is fully predicted, that is, both inputs are equal, the PE neurons remain at their baseline. Mismatches can come in two flavors: either the predicted sensory input is larger than the actual sensory input (‘Overprediction’, in (6) this phase is referred to as ‘mismatch phase’), or smaller than the actual sensory input (‘Underprediction’, in (6) this phase is referred to as ‘playback phase’). While nPE neurons only increase their firing rate relative to baseline when the sensory input is overpredicted, pPE neurons increase their activity when the sensory input is underpredicted.

### A multi-pathway E/I balance in negative and positive prediction-error neurons

To investigate the conditions under which both PE neuron types co-exist, we first made use of a simplified mean-field analysis of the full neural network, for which the dynamics of each neuron type or compartment are represented by a linear equation. This allowed us to perform an exhaustive mathematical analysis of the steady-state responses of all neuron types (see Supporting Information for details).

We found that in a network with inhibitory PV, SOM and VIP neurons, both nPE and pPE neurons can co-exist when the interneuron network establishes a balance of excitation and inhibition (see also 23, for nPE circuits). An informative example of such a balance is a network in which SOM neurons receive sensory inputs and VIP neurons receive a prediction thereof (Fig. 1B-E). Both nPE and pPE neurons receive at their soma the same amount of excitatory and inhibitory inputs when the network is stimulated with fully predicted sensory stimuli. For nPE neurons, this balance is preserved for stimuli larger than predicted, and temporarily broken in favor of excitation for stimuli smaller than predicted. In contrast, for pPE neurons, the E/I balance is preserved for stimuli smaller than predicted, and temporarily broken for stimuli larger than predicted (Fig. 1B).

Importantly, our analysis shows that this E/I balance is not only a balance in the total inputs onto PE neurons but also the pathways those inputs can take through the circuit (Fig. S1, and Methods for details). To show this, we computed for each neuron type/compartment the sum of all pathways that originate from this node and ended either at the soma or the dendrites of PE neurons. The contributions for all neuron types/compartments, separated into net excitatory and inhibitory pathways, reveal an E/I balance (Fig. 1C, and Fig. S2 B). In a recurrent neural network with different types of inhibitory neurons, the individual pathways comprise a number of excitatory, inhibitory, dis-inhibitory and dis-disinhibitory connections (also see 23). A consequence of the balanced pathways is that the ability of PE neurons to remain at their baseline activity is independent of the particular stimulus strength, provided the neuronal input-output transfer functions are sufficiently linear.

A mean-field network with a connectivity satisfying the constraint of a compartment-specific E/I balance shows both nPE and pPE neurons (Fig. 1D). The activity of PV, SOM and VIP neurons varies between the different phases, reflecting the underlying connectivity required to achieve a multi-pathway E/I balance, and the different inputs onto the interneuron types. In our PE circuit, mismatch responses are the result of an excess of excitation at the soma of one PE neuron type and a withdrawal of inhibition at the dendrites of the other PE neuron type, respectively (Fig. S2B). In line with mismatch neuron responses in the primary visual cortex (6), the simulated mismatch responses increase with the difference between actual and predicted sensory inputs (Fig. 1E).

Our analysis revealed that a circuit with only one source of somatic inhibition was not sufficient to give rise to both nPE and pPE neurons. This can be intuitively understood by the following reasoning. The dendrites are balanced or inhibited for one of the two mismatch phases (Fig. S2), and, hence, do not contribute to the somatic activity. As a result, any change of activity in PE neurons must be a consequence of changes in the somatic inputs. The PE neuron type increasing the activity during that mismatch phase requires the currents flowing through the interneuron circuit to be unbalanced. However, the PE neuron type that remains at its baseline during that mismatch phase requires the currents flowing through the interneuron circuit to be balanced (see derivations in Supporting Information). These two conditions can not be satisfied in an interneuron circuit in which the currents are directed through one soma-targeting interneuron population only. In our network, somatic inhibition is provided by PV neurons. To create two types of somatic inhibition, we, therefore, subdivided PV neurons into two groups, one receiving sensory inputs, the other one receiving a prediction thereof. Given the vast diversity of feedforward and feedback inputs to PV neurons reported experimentally (38, 43, 44), this assumption is plausible for real biological systems. However, besides PV neurons, other soma-targeting interneurons (11) can contribute to establishing an E/I balance, so that the division of PV neurons into sub-populations is not a strict requirement. The results are robust to the input distribution onto SOM and VIP neurons (Fig. S2). Changing their inputs only affects the pathway strengths that are required to achieve an E/I balance. However, we find that for mismatch responses to develop, either SOM neurons, VIP neurons or both must receive the actual sensory input (Fig. S2).

In summary, our analysis shows that in a simplified mean-field network, PE neurons with arbitrary baseline activity require a multi-pathway E/I balance. Mismatch responses are the consequence of a temporary imbalance of excitation and inhibition, caused either by an excess of somatic excitation or a withdrawal of dendritic inhibition. Moreover, the co-existence of nPE and pPE neurons requires at least two distinct sources of somatic inhibition, as well as dendritic inhibition that is also driven by sensory inputs.

### Prediction-error neurons are robust to network manipulations

The background inputs to all neurons were taken to be fixed in our simulations. However, neurons are constantly bombarded with non-stationary local and long-range synaptic inputs, and regulated by neuromodulators like acetylcholine or dopamine. Moreover, the activity of both excitatory and inhibitory neurons is modulated by behavioral states, and highly context-dependent (45). This naturally leads to the question of whether nPE and pPE neurons are robust to network perturbations. If nPE and pPE neurons are sensitive to small changes in the inputs, they would need to be reconfigured constantly. To study the network’s ability to withstand perturbations, we individually injected additional inhibitory or excitatory inputs to the neurons of our PE circuits (Fig. 2A). In our analysis, we focused on moderate perturbation strengths to ensure that none of the neuron types is silenced.

**Figure 2.**
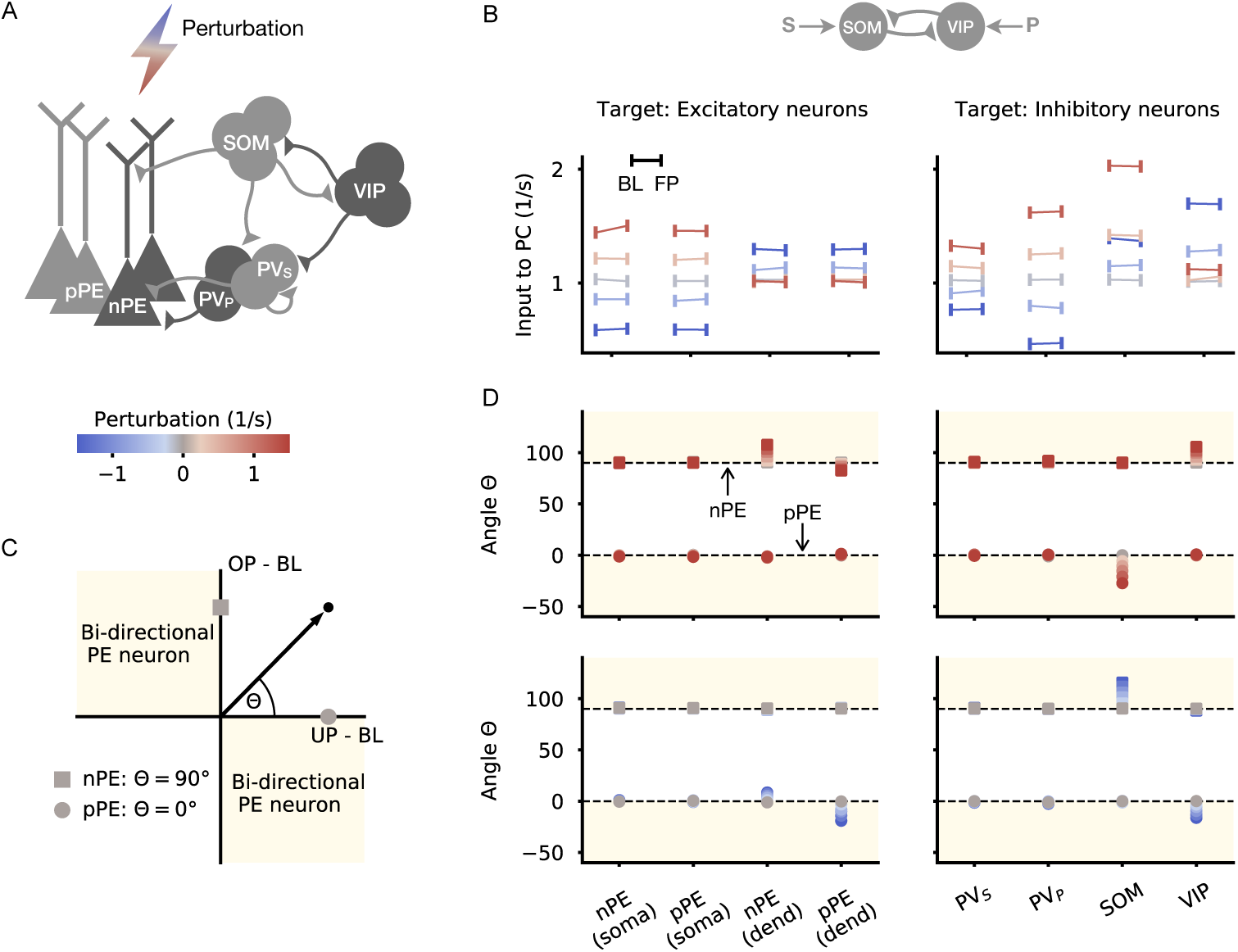
PE neurons are robust to moderate network perturbations. **(A)** Each neuron type/compartment of a mean-field network with nPE and pPE neurons is perturbed with an additional inhibitory or excitatory input. Same circuit as in Fig. 1B-E. SOM neurons receive the actual sensory input while VIP neurons receive a prediction thereof. **(B)** Total input into PE neurons during the absence of sensory stimuli (baseline, BL) and for fully predicted sensory stimuli (FP) for different perturbation strengths and different perturbation targets (Left: compartments of PCs, Right: inhibitory neurons). Total input in both phases are almost equal as a result of the established E/I balance. Gray: no perturbation. **(C)** Illustration of nPE (square) and pPE (circle) neurons in the input space. Input space is defined by the total input to PCs for overpredicted (OP) and underpredicted (UP) sensory stimuli. nPE neurons lie on the positive part of the y-axis, while pPE neurons lie on the positive part of the x-axis. Beige areas denote bi-directional PE neurons. Θ defines the angle in the input space. **(D)** Θ for different perturbation strengths (top: excitatory, bottom: inhibitory) and different perturbation targets (Left: compartments of PCs, Right: inhibitory neurons). Perturbations have minor effects on nPE and pPE neurons, especially when baseline firing rates of PCs are low.

A unifying hallmark of both nPE and pPE neurons is that they remain at their baseline for fully predicted stimuli. Hence, the total input to PE neurons for anticipated stimuli must be equal to the total input in the absence of sensory stimuli. Both excitatory and inhibitory network perturbations can change the inputs to PE neurons in the baseline phase (Fig. 2B and Fig. S3). However, the total inputs for fully predicted sensory stimuli change to the same extent (Fig. 2B and Fig. S3), leading to no significant changes in activity relative to baseline.

In the next step, we wondered how nPE and pPE neurons change their responses to unexpected mismatches when these mismatches are accompanied by neuron-specific perturbations. For each perturbation target and strength, we plotted the total input for over- and underpredicted sensory stimuli. In this depiction, nPE neurons lie on the positive part of the y-axis, while pPE neurons lie on the positive part of the x-axis (Fig. 2C). The second and fourth quadrants denote the range of bi-directional PE neurons that either increase activity for sensory inputs smaller than predicted and decrease activity for sensory inputs larger than predicted, or vice versa. We quantify perturbation-induced changes of PE neuron activity by the angle Θ in this input-space (Θ = 90: nPE neurons, Θ = 0: pPE neurons).

Perturbations that targeted the soma of nPE and pPE neurons, either directly or indirectly through PV neurons, have no or only comparatively small effects on the responses upon unexpected mismatches (Fig. 2D). Perturbations that targeted the dendrites, either directly or indirectly through SOM and VIP neurons, can have salient effects for some of the perturbation strengths tested. In those cases, uni-directional PE neurons mostly transition into bi-directional PE neurons. However, when excitatory neurons have very low (close to zero) baseline activities, negative deviations from the baseline are bounded from below, and hence, are undetectable.

Altogether, these perturbation experiments show that once an E/I balance has been established and gives rise to nPE and pPE neurons, these neurons are robust to moderate network manipulations, and, hence, do not need to be reconfigured. Moreover, our simulations indicate that perturbations of either the dendrites of PCs, inhibitory SOM or VIP neurons can modulate the mismatch responses of PE neurons.

### Negative and positive prediction-error neurons develop through inhibitory plasticity with a low homeostatic target rate

It has been shown that mismatch neurons in the primary visual cortex are experience-dependent and require visuomotor coupling to develop normally (6). This suggests that PE neurons are formed through learning. In a model of rodent V1, nPE and pPE neurons were learned separately by means of local, homeostatic inhibitory plasticity (23). It is, however, not resolved how this can be generalized to learning nPE and pPE neurons in the same recurrent network simultaneously.

The homeostatic inhibitory plasticity used in (23) keeps the PCs at a target rate by adjusting the connectivity such that to establish an E/I balance for all inputs the network is exposed to. A direct consequence is that nPE neurons may form when the network is exposed to stimuli that are smaller than or equal to the prediction. Likewise, pPE neurons may develop when the network is exposed to sensory stimuli that are larger than or equal to the prediction. The increase of activity in nPE neurons for overpredicted stimuli and in pPE neurons for underpredicted stimuli is, hence, a result of the network never experiencing the respective mismatch during training. This clearly shows a dilemma: to learn nPE and pPE neurons in the same network, it must be exposed to fully predicted as well as over- and underpredicted stimuli. The plasticity rule, however, will keep the excitatory neurons at a target rate throughout, so that neither nPE nor pPE neurons could emerge.

We found that a simple solution is to train the network only with phases of fully predicted sensory inputs, and set the homeostatic target rate of excitatory neurons to zero. This way, the neurons learn to remain at their baseline rate for fully predicted stimuli. At the same time, any excess of somatic inhibition for over- and underpredicted stimuli is not reflected in the firing rates because it is bounded by zero. To show that in this way both nPE and pPE neurons can develop, we made the inhibitory connections onto the soma and dendrite of PCs as well as the inhibitory connections from SOM and VIP neurons onto PV neurons subject to experience-dependent plasticity (Fig. 3A). While the synapses onto PCs follow an inhibitory plasticity rule akin to (46), the inhibitory synapses onto PV neurons follow a local approximation of the backpropagation of error rule (47) that relies on information forwarded to the PV neurons from the PCs (see also 23, 48).

**Figure 3.**
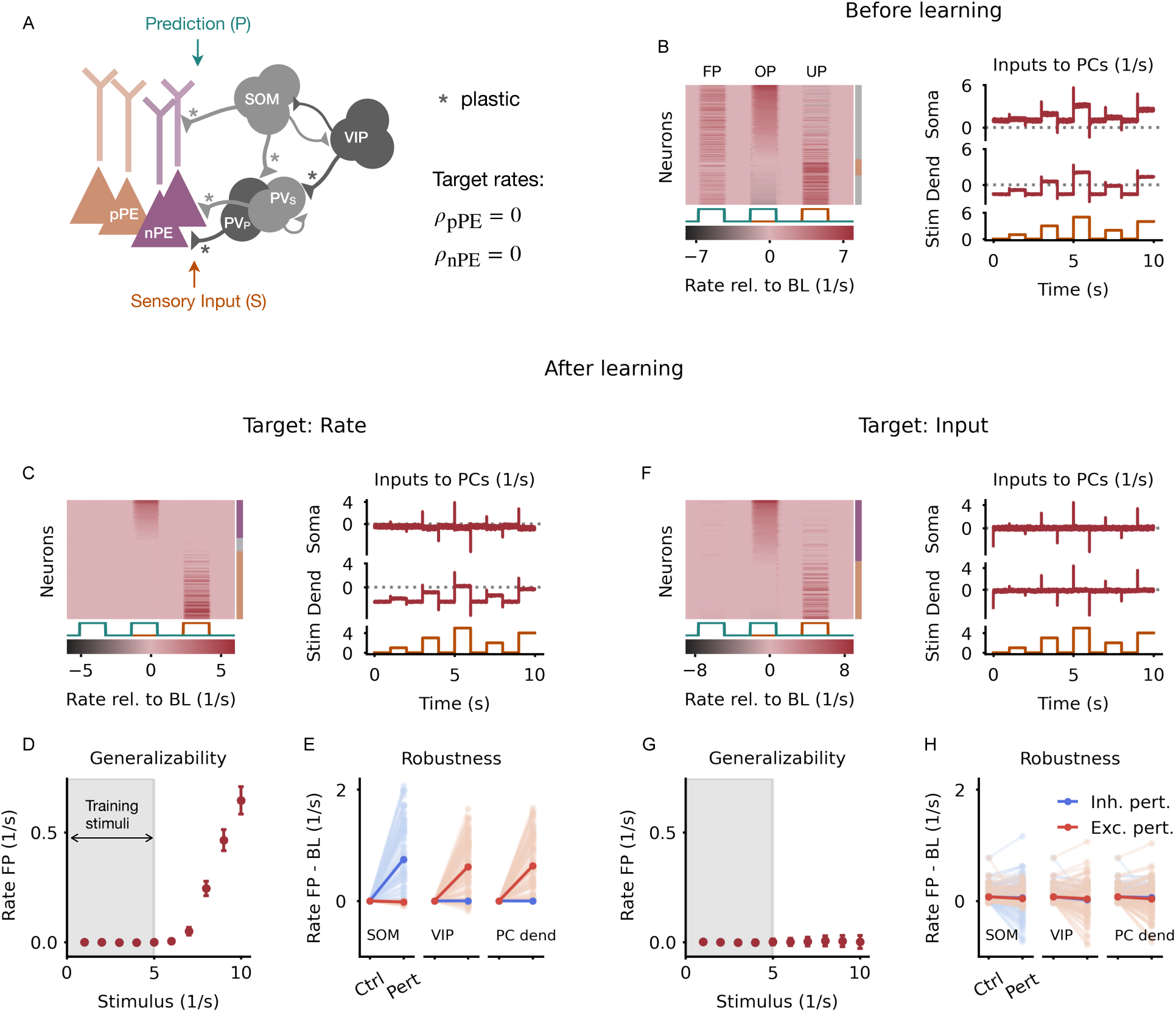
nPE and pPE neurons develop through inhibitory plasticity with a low homeostatic target rate. **(A)** Heterogeneous network model with excitatory PCs and inhibitory PV, SOM and VIP neurons. All PCs receive actual sensory input at the somatic compartment and a prediction thereof at the dendritic compartment. 50% of the PV neurons, 70% of the SOM neurons and 30% of the VIP neurons receive the sensory stimuli. The remaining cells receive the prediction. Connections marked with an asterisk undergo experience-dependent plasticity. Target rates for PCs are set to zero. **(B)** Responses of and inputs to PCs before learning. Left: Responses relative to baseline of all PCs for fully predicted (FP), overpredicted (OP) and underpredicted (UP) stimuli, sorted by amplitude of mismatch response in OP. Almost none of the PCs are classified as PE neurons summarized by the bar to the right (gray: no PE neuron, purple: nPE neuron, orange: pPE neuron). Right: Mean input into both soma and dendrites of PCs for fully predicted stimuli. Inputs are not balanced. **(C)** Same as in B but after learning with an inhibitory plasticity rule that establishes a zero target rate in PCs. Left: Most of the PCs are either nPE (purple) or pPE (orange) neurons (indicated by the colored bar to the right). Right: Mean input into both soma and dendrites of PCs for fully predicted stimuli are not balanced. **(D)** Median and SEM of PC responses for fully predicted sensory stimuli. Gray area indicates the range of stimuli used during learning. Sensory stimuli that are larger than the training stimuli evoke neuron responses. **(E)** Inhibitory (blue) and excitatory (red) perturbations can cause the PE neurons to deviate from their baseline activity. Light colors denote single neurons, dark colors denote the population average. Ctrl: Control, Pert: Perturbation. **(F-H)** Same as C-E but with an inhibitory plasticity rule that establishes a target for the total input to PCs (target: zero). **(F)** Left: Most of the PCs are either nPE (purple) or pPE (orange) neurons (indicated by the colored bar to the right). Right: Mean input into both soma and dendrites of PCs for fully predicted stimuli are balanced. **(G)** Sensory stimuli that are larger than the training stimuli evoke only minor neuron responses. The PE neurons can generalize beyond the training stimuli. **(H)** PE neurons are robust to inhibitory and excitatory perturbations after learning.

Before learning, the network was randomly initialized with a connectivity motivated by experimental studies (12, 13, 30–35), leading to PCs that show deviations from baseline in all phases (Fig. 3B, left). The excitatory and inhibitory inputs at both soma and dendrites of PCs are unbalanced for fully predicted stimuli (Fig. 3B, right). Only very few neurons could therefore be classified as PE neurons. The total inputs to PCs for over- and underpredicted stimuli were negatively correlated, and showed a near balance of top-down predictions and bottom-up sensory inputs (49) (Fig. S4A). During learning, the inhibitory synapses collectively adjusted their efficacy to keep the PCs at their target rate for perfectly predicted sensory inputs (Fig. S5A). At the end of learning, the majority of PCs showed response patterns akin to nPE or pPE neurons (Fig. 3C, left). The balance of top-down predictions and bottom-up sensory inputs was preserved after learning (Fig. S4B). However, the excitatory and inhibitory inputs at both soma and dendrites of PCs are not perfectly balanced (Fig. 3C, right). That is a consequence of the target rate being equal to the neurons’ rectification threshold. The objective function (*r*_PC_ = *r*_target_ = 0) is already satisfied when the total input to PCs is less or equal to zero for all stimuli presented during training. That is, the network does not necessarily strive for an E/I balance. While this does not hamper the formation of nPE and pPE neurons per se, it compromises some of the properties of PE neurons that emerge from this balance. On the one hand, the ability to generalize beyond the training stimuli does not hold (Fig. 3D). On the other hand, the network is less robust to perturbations (Fig. 3E), which means that PE neurons would have to be relearned continuously.

We, therefore, modified our plasticity rules by introducing a target for the total input to PCs, instead of a target for their firing rate. This allows the synaptic weights to adjust to both positive and negative deviations from the target, and forces the network to establish a balance of excitation and inhibition. After learning, excitatory and inhibitory inputs to both soma and dendrites are balanced on a stimulus-by-stimulus basis (Fig. 3F, right). As before, almost all PCs developed into nPE or pPE neurons (Fig. 3F, left), and the balance of top-down predictions and bottom-up sensory inputs is preserved. (Fig. S4C). Finally, nPE and pPE neurons generalize beyond the training stimuli (Fig. 3G), and PE neurons are more robust to moderate network perturbations (Fig. 3H).

This shows that networks with inhibitory plasticity can give rise to both nPE and pPE neurons when the homeostatic target rate is zero. Moreover, when the plasticity acts to establish a target for the total input to excitatory neurons, the PE neurons’ ability to withstand network perturbations and generalize is improved.

### Mismatch responses are determined by initial connectivity and inputs onto the interneurons

Training a network solely with fully predicted sensory inputs (Fig. 3) does not constrain the neuron responses to mismatches. Theoretically, the PC responses to over- and underpredicted stimuli can be classified into four cases. When PCs are silent in the absence of sensory inputs, these neurons would be either nPE neurons, pPE neurons, silent neurons, or neurons that indicate mismatches independent of the valence. We therefore wondered which network properties determine the ratio between nPE and pPE neurons.

We find that a key factor for whether PCs develop into nPE or pPE neurons is the initial connectivity before learning. A mean-field analysis of homogeneous PE circuits shows that nPE and pPE neurons take up distinct parameter ranges in the weight space (Fig. 4A, top). When the initial connectivity is closer to the nPE-manifold, PCs tend to develop into nPE neurons (Fig. 4B, left). Likewise, when the initial connectivity is closer to the pPE-manifold, PCs tend to develop into pPE neurons (Fig. 4B, middle). Hence, for networks to give rise to both types of PE neurons, the initial connectivity should comprise sufficiently large parameter spaces, or regions close to both manifolds (Fig. 4B, right).

**Figure 4.**
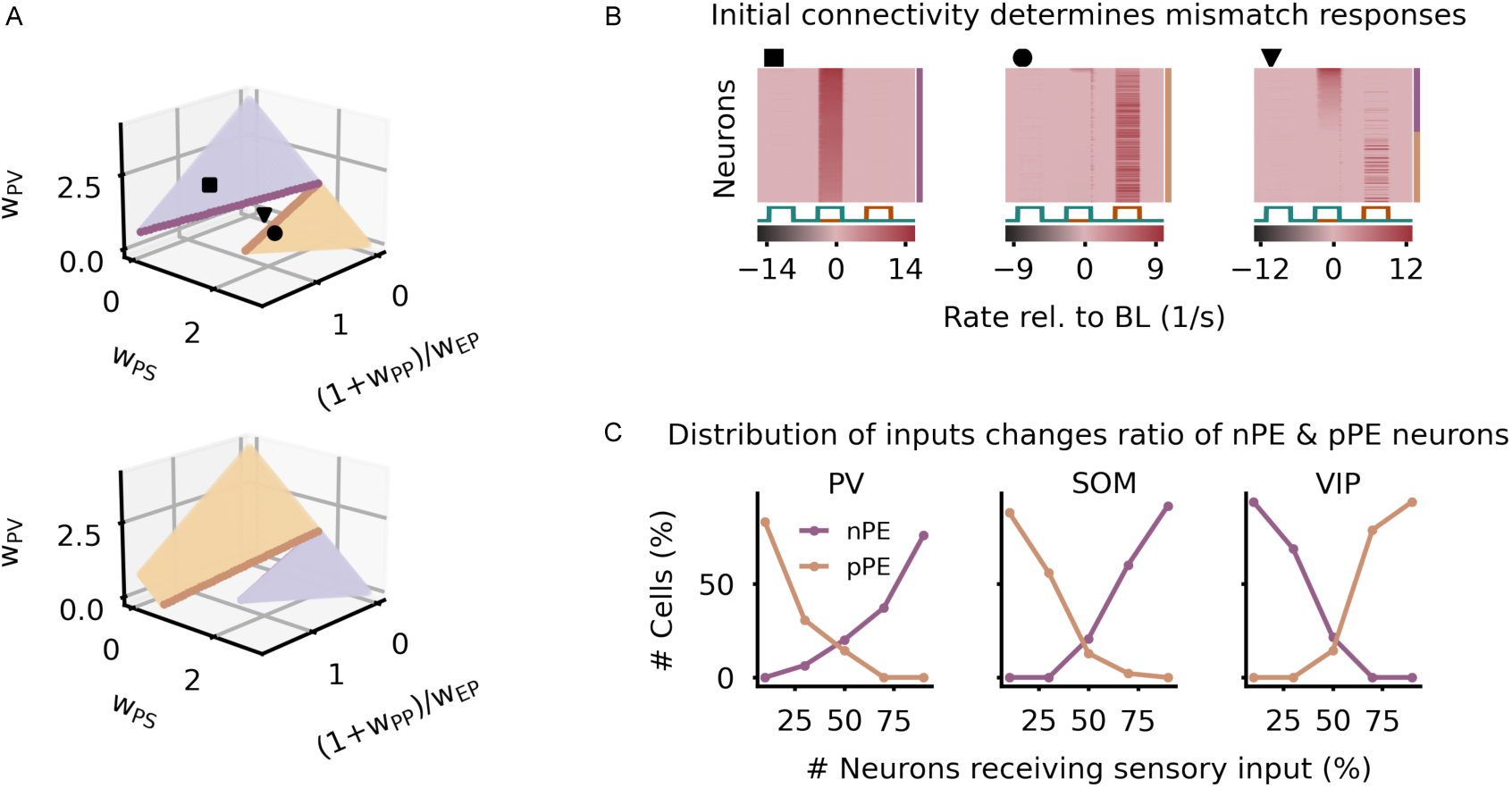
Initial connectivity and distribution of inputs onto interneurons determine mismatch responses of PE neurons. **(A)** nPE (purple) and pPE neuron (orange) manifolds in the parameter space. Manifolds derived from a mean-field analysis of separate nPE or pPE circuits. PV neurons receive both actual and predicted sensory inputs. SOM neurons receive sensory inputs, while VIP neurons receive a prediction thereof (top), or vice versa (bottom). Markers denote different initial states. For the sake of visualization, (1 + *w*_PP_)*/w*_EP_ was increased by 0.5. *w*_EP_: weight from PV neurons onto soma of PCs, *w*_PP_: recurrent inhibition between PV neurons, *w*_PS_: weight from SOM neurons onto PV neurons, *w*_PV_: weight from VIP neurons onto PV neurons. **(B)** For three different initial weight configurations (A), the network forms nPE neurons (left), pPE neurons (middle), or both (right). **(C)** The number of PV neurons (left), SOM neurons (middle), or VIP neurons (right) that receive the actual sensory input is increased. The ratio of nPE and pPE neurons changes with the distribution of actual and predicted sensory inputs onto the interneurons.

Another crucial factor for the ratio of nPE and pPE neurons is the distribution of actual and predicted sensory inputs onto the interneurons. Changing the distribution of those inputs onto PV, SOM and VIP neurons changes the parameter ranges for nPE and pPE neurons (Fig. 4A, bottom). When the majority of PV or SOM neurons receive the actual sensory input, PCs tend to develop into nPE neurons (Fig. 4C). Likewise, by increasing the number of VIP neurons that receive a prediction of the expected sensory input, most of the excitatory neurons become nPE neurons after training. If these ratios are inverted, the PE circuit is biased towards pPE neurons.

Finally, the predictability of sensory stimuli during training can bias the formation of PE neurons. Given that neuronal networks receive substantial noise due to, for instance, the random nature of synaptic transmission, synapse failures or channel noise (50), sensory inputs and predictions thereof will rarely be perfectly equal. As a consequence, networks are not only exposed to perfectly predicted sensory stimuli but also to small mismatches between them. When the transmission of the predicted sensory input becomes less reliable, the number of pPE neurons strongly decreases up to a point where no pPE neurons are formed during learning (Fig. S6A). Similarly, when the transmission of the actual sensory input becomes less reliable, the number of nPE neurons strongly decreases up to a point where no nPE neurons are formed (Fig. S6B). While both the numbers of nPE and pPE neurons decrease equally when the network is exposed to noisy stimuli, PE neurons can still form. This reflects the network’s ability to tolerate positive and negative deviations between actual and predicted sensory inputs when both phases are presented equally (Fig. S6C).

Altogether, this shows that the initial connectivity, distribution of actual and predicted sensory inputs onto interneurons, and stimulus predictability during development determine which PE neurons emerge, and that the formation of nPE and pPE neurons is robust with respect to variations in these parameters.

### Prediction-error neurons bias unpredictable percepts towards the mean of the stimulus statistic

Previous experiences shape perception and behavior. A salient example of experience-dependent biased perception is *bias towards the mean* (aka contraction bias). A stimulus drawn from a random distribution is perceived larger when it is smaller than the mean of the stimulus distribution, and perceived smaller when it is larger than the mean. This well-known phenomenon has been described centuries ago (51, 52), reproduced many times in tasks that involve the reproduction of a perceived variable (24–29), and attributed to Bayesian computation in which a system integrates information about prior stimulus statistics (25, 26). We speculated PE neurons can support biased perception.

To this end, we simulated two connected subnetworks, the neurons of which are only responsive to a range of stimuli. Each subnetwork consists of a PE circuit as studied before, connected with a representation neuron and an attractor network (Fig. 5A). The attractor network consists of two memory neurons modeled as perfect integrators (that is, line attractors) and two prediction neurons. Each memory neuron projects to the associated prediction neuron of that subnetwork. The prediction neurons are mutually connected via inhibitory synapses (53), forming two fixed points. Neurons of the attractor network receive inputs from both nPE and pPE neurons of their associated subnetwork. While nPE neurons inhibit the attractor neurons (for instance, through inhibitory interneurons not explicitly modeled here), pPE neurons excite them (motivated by 3). The representation neurons, whose activity represents the perceived stimulus, not only receive the sensory stimulus itself but are also connected to both nPE and pPE neurons of their respective subnetwork, with reversed connectivity (Fig. 5A). For simplicity, we assume that during the presentation of random stimuli, the PE circuits are not updated significantly, for instance because the learning rate is small compared to the changes in activity, or even simply because the effect of positive and negative mismatch phases is balanced (Fig. S6C). The strength of sensory stimuli is drawn from two uniform distributions, one ranging from 1 to 5 *s*^−1^ (associated with the first subnetwork), and the other one ranging from 5 to 9 *s*^−1^ (associated with the second subnetwork). In addition, the distribution is indicated to the network by a cue signal delivered to the prediction neurons.

**Figure 5.**
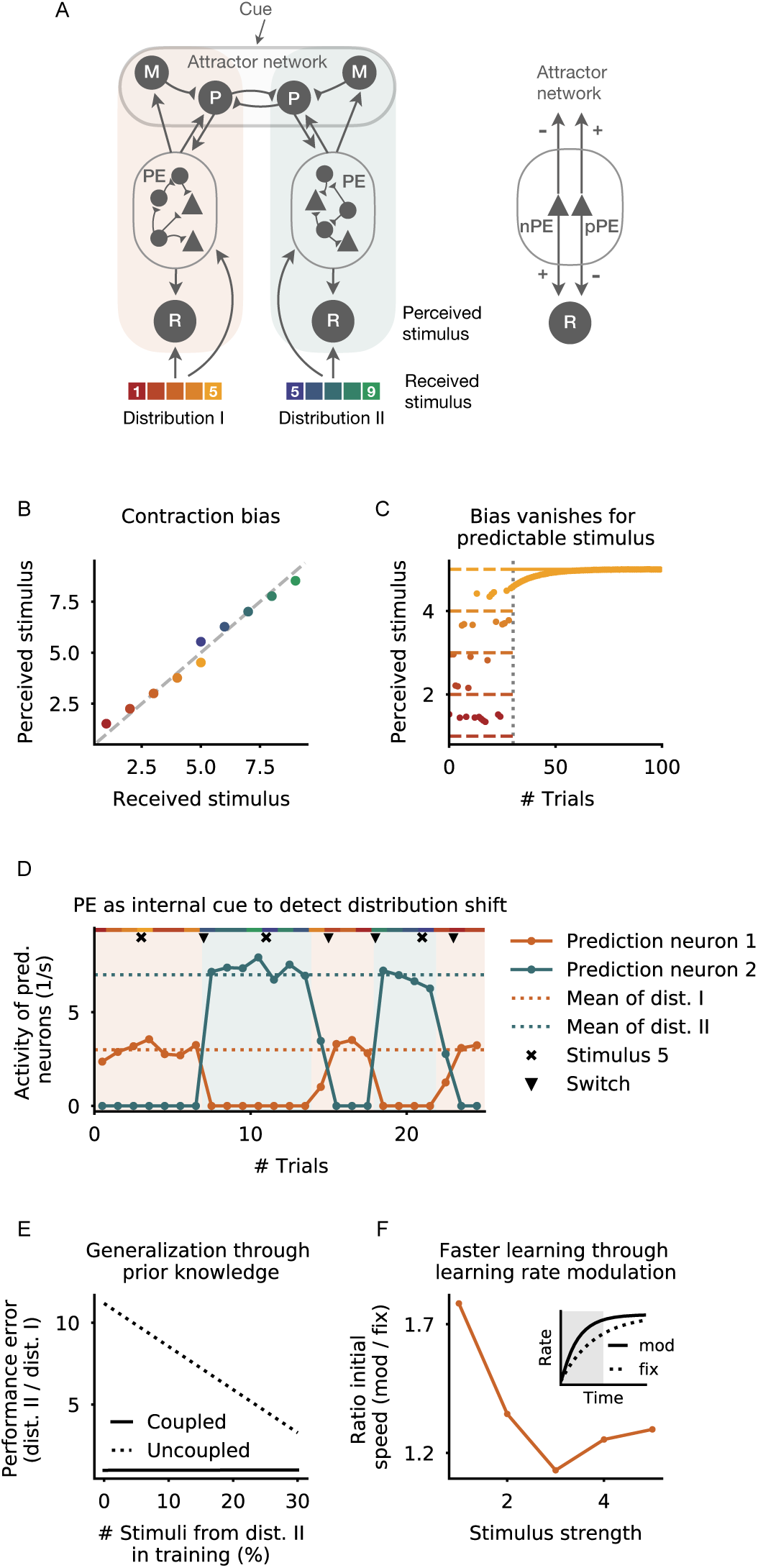
The role of PE neurons in biased perception. **(A)** Left: Attractor-memory network with PE neurons. The network consists of two subnetworks, the neurons of which are only responsive to a subset of stimuli. Each subnetwork comprises a representation neuron (R) and a PE circuit. Both R and PE neurons receive sensory stimuli of either of two uniform distributions. The PE circuit is connected to both the representation neuron (R) and an attractor network that comprises memory neurons (M) and prediction neurons (P). The two prediction neurons are mutually connected via inhibitory synapses, and receive excitatory input from the memory neuron of their respective subnetwork. Right: nPE and pPE neurons connect to M, P and R neurons with opposing sign. **(B)** The PE neurons establish a contraction bias for both distributions. A stimulus that is smaller than the distribution mean is perceived stronger, while a stimulus that is larger than the distribution mean is perceived weaker. **(C)** The response of the representation neuron becomes unbiased after the transition (dotted vertical line) from a uniform distribution to a binary distribution because the stimulus becomes predictable. **(D)** The network does not receive a cue signal indicating the distribution from which the stimuli are drawn. After an uncued switch from one distribution to another, the former inactive prediction neuron becomes active and the former active prediction neuron becomes inactive (network switching denoted by triangle). This is the result of the PE neurons and the mutual inhibition between both prediction neurons. Stimulus present in both distributions does not evoke a switch (denoted by x). Shaded areas denote the distributions form which the stimuli are drawn. **(E)** Both distributions equally change from uniform to binary (to maximal values of former uniform distributions). A network in which the PE neurons are equally coupled to both memory neurons (solid line) shows the same performance error for both distributions independent of the training set composition. A network in which the PE neurons are only coupled to the memory neuron of their respective subnetwork (dashed line) shows a larger error for the distribution that is underrepresented during training. **(F)** Speed of learning (defined as the averaged change of activity in the first 50 ms; gray area in inset) is increased when PE neuron activity modulates the learning rate based on the degree of the stimulus’ unpredictability, compared to a fixed learning rate (solid vs. dashed line in inset).

#### Contraction bias for unpredictable and predictable stimuli

In our network, the memory neurons hold the mean of the past stimuli presented. When sensory stimuli are drawn from a unimodal distribution, the activity of the memory neuron approaches the mean of that distribution. Deviations from this mean are controlled by nPE and pPE neurons. When a prediction neuron is active while the other one is silent, its steady-state activity is, hence, also given by the mean of the distribution, but modulated by the momentary input of the PE neurons. Sensory inputs that are smaller than the activity of the prediction neuron activate nPE neurons. Because nPE neurons excite the representation neuron of that subnetwork, the perceived stimulus is larger than the received stimulus. In contrast, sensory inputs that are larger than the activity of the prediction neuron activate pPE neurons (Fig. 5B). Because we assume that the net effect of pPE neurons on representation neurons is negative, the perceived stimulus is smaller than the received stimulus. This effect is particularly pronounced for the stimulus present in both distributions (i.e., 5 *s*^−1^).

The preceding results suggest that the bias is a consequence of the unpredictability of the stimulus. Hence, when the sensory input becomes predictable, the bias should eventually vanish. To test this, after some trials with random stimuli, we always presented the same sensory input. The activity of the PE neurons slowly changes the memory neuron until it represents the actual stimulus. As a result, over time, the prediction neuron itself represents the sensory input, a consequence being that the PE neurons become silent. Hence, the bias vanishes, and the perceived stimulus equals the received stimulus (Fig. 5C).

#### Prediction-error neurons may act as internal cue

While the distribution from which the stimulus is drawn had been cued to the network so far, very often changes in the environment or underlying tasks occur spontaneously without clues. We figured that PE neurons may act as internal cues that support a switch between attractors when stimuli are suddenly drawn from the other distribution. Immediately after the network experiences an unexpected switch, the prediction neuron that had been active in the previous trials remains active. However, after some time, the PE neurons of that subnetwork suppress the activity of the prediction neuron. At the same time, the PE neurons of the other subnetwork excite the corresponding prediction neuron. Together with the mutual inhibition between prediction neurons, the wrong expectation is eventually corrected successfully (Fig. 5D). Importantly, stimuli that are present in both distributions are not sufficient to cause a switch (Fig. 5D, see stimuli denoted by x). Altogether, this shows that PE neuron activity may underlie fast adaptation to unexpected situations by forcing the attractor network to switch between fixed points.

#### Generalization through prior knowledge

The activity of PE neurons might also support generalization across environments or tasks by making use of prior knowledge encoded in the connectivity between the PE circuit and the attractor network. To illustrate this, let us assume that any change of one distribution will equally affect the other distribution. In our network, this can be implemented by cross-coupling the PE neurons of one subnetwork with the memory neuron of the other subnetwork. As a result, any changes in input statistics will equally affect both memory neurons, even when the network is only exposed to samples of one distribution. To confirm this, we changed both distributions from uniform to binary. After the transition, each distribution only comprises the maximal stimulus of the former uniform range of stimuli. We then trained the network briefly with the new input statistics. While networks without cross-coupling show larger performance errors for the distribution that was underrepresented during training, networks with cross-coupling show the same test error for both distributions independent of the training set composition (Fig. 5E). Hence, PE neurons can support generalization by modulating memory neurons of all subnetworks.

#### Faster learning through modulation of learning rates

Finally, PE neurons could also facilitate learning by adjusting learning rates based on the degree of predictability of sensory stimuli. In such a scenario, unexpected stimuli would increase the learning rate, leading to faster adaptation of synaptic connections. On the neuronal level, this could be achieved by PE neurons targeting the distal locations of basal dendrites, while the bottom-up sensory inputs arrive at the proximal locations of the basal dendrites. The distal synapses may then only elicit NMDA spikes that cause the proximal synapses to strengthen (54). To illustrate this, we repeatedly stimulate subnetwork 1 with the same stimulus and update the synaptic weight connecting the stimulus with the representation neuron. To quantify the speed-up in learning, we compute the rate of change in the activity of the representation neuron at the beginning of training for both a learning rate that is fixed and for one that is modulated by the activity of PE neurons (Fig. 5F). As expected, the speed-up is larger for stimuli that deviate more from the distribution mean. This shows that modulating learning rates by the degree of unpredictability (or surprise) of an event can underpin fast learning.

## Discussion

We showed that, in a mean-field network, nPE and pPE neurons with arbitrary baseline activity co-exist when an E/I balance is established at the soma and dendrite of PCs for fully predicted sensory stimuli. While for nPE neurons the balance at the soma is preserved for underpredicted stimuli, excitation and inhibition cancels for overpredicted stimuli in pPE neurons (Figs. 1 and S2). Moreover, we reveal that the balance of inhibitory and excitatory inputs emerges from balanced pathways (Figs. 1, S1 and S2) that include, among others, excitatory, inhibitory, disinhibitory, and dis-disinhibitory pathways (Fig. S1). Based on a mathematical analysis, we showed that for both PE neurons to co-exist, somatic inhibition must come in two distinct variants (Supporting Information). In addition, the interneurons providing dendritic inhibition should be driven by actual sensory inputs. Otherwise, mismatch responses vanish (Fig. S2).

For a heterogeneous plastic network, we demonstrated that nPE and pPE neurons can simultaneously develop by inhibitory plasticity with a homeostatic target firing rate that keeps PCs silent in the absence of stimuli (Figs. 3 and 4). When the plasticity rule is modified to process mismatches from a target input, PE neurons generalize beyond the range of sensory stimuli seen during learning, and are robust to network perturbations (Figs. 3 and 2). Finally, we showed that an attractor-memory network with a PE circuit can reproduce the contraction bias for unpredictable stimuli (Fig. 5). We demonstrated, by means of the example of biased perception, that PE neurons can act as an internal cue that indicates unannounced switches between stimulus distributions. Moreover, we illustrated that PE neurons may underpin generalization across stimulus statistics, and can support faster learning (Fig. 5).

Our work makes a number of predictions that could be tested experimentally. (1) PE neurons arising from balanced pathways are robust to network perturbations, that is, they remain at baseline for fully predicted sensory inputs. (2) If PE neurons change upon perturbation, it mainly affects their responses to unpredicted stimuli. Those changes primarily occur for direct or indirect (through SOM and VIP neurons) perturbations of the dendrites, and usually lead the former unidirectional PE neurons to act as bidirectional PE neurons. These predictions can be tested by optogenetically or pharmacogenetically manipulating neuron types/compartments. (3) Both nPE and pPE neurons are hidden bidirectional PE neurons because of their low baseline firing rate. This can be directly tested by elevating their baseline activity through external excitatory stimulation because additional input should not affect the PE neurons’ ability to remain at their baseline activity for fully predicted sensory stimuli (see above). (4) PE neurons generalize beyond the stimuli used during learning. By carefully designing experiments that restrict learning to a subset of stimuli, the developing PE neurons can be tested for their ability to generalize. (5) PE neurons underlie contraction bias. It means the observed bias should vanish for targeted silencing of PE neurons. Moreover, if only one of the two PE neuron types is silenced, the bias would only occur for one side of the stimulus distribution. Although still challenging, by employing recent technological advances, for instance, multiphoton holographic optogenetics (55) and neuron tagging based on activity-dependent promoters (56, 57), targeted manipulation of nPE and pPE neurons may be in reach soon.

While, in the present study, we have focused on a canonical interneuron circuit that has been described widely (see, for instance, 12, 13, 32), our mathematical analysis can be straightforwardly extended to an arbitrary number of interneurons. In the motif with PV, SOM and VIP interneurons, somatic inhibition is provided by PV neurons, while dendritic inhibition is provided by the SOM-VIP circuit. nPE and pPE neurons can also emerge without VIP neurons (see discussion and Supporting Information in 23). However, VIP neurons in our network (i) contribute to amplify mismatch responses (6, 22), and (ii) allow SOM neurons to receive both the actual and the predicted sensory inputs, respectively (Fig. S2). Input heterogeneity is in line with studies showing that interneurons receive both feedforward and feedback inputs (38, 43, 44).

In our simulations, the external input onto the dendrites during learning is chosen such that the dendrite is silent in the absence of any sensory input. Further sources of dendritic inhibition in the form of other dendrite-targeting interneuron types may support establishing a target activity in the dendrites, and the formation of prediction errors. For example, a number of interneuron types are located in layer 1 (58, 59), the major target for feedback connections (37, 60, 61). NDNF neurons, for instance, have been shown to inhibit the apical dendrites located in the superficial layers (62). These neurons could, hence, be a promising candidate in shaping dendritic prediction errors.

While we have modeled the interneurons as point neurons, the PCs were modeled as two coupled point compartments that represent soma and dendrites of PCs. We assume that a rectifying nonlinearity at the dendrite decouples the apical tuft of PCs from their soma when dendritic inhibition exceeds excitation. Many biological mechanisms are, however, absent from our model. First, the dendritic tree was simplified to one compartment, neglecting the potential influence of dendritic branches to process distinct information (20). Second, we do not explicitly model calcium spike-like events at the dendrites (63, 64). However, we have shown previously that nPE neurons emerged in the same way in a model with the potential to produce calcium spike-like events (23). Hence, we expect our results to generalize to more elaborated dendritic structures.

In our analysis, we assumed that the apical dendrites of PCs are balanced for fully predicted sensory stimuli, and can form a prediction error for unexpected inputs, which is then transmitted to the soma of PCs (see also, 65). Depending on the distribution of actual and predicted sensory inputs onto SOM and VIP neurons, and depending on whether the PC forms an nPE or pPE neuron, their dendrites can either be balanced, over-inhibited, or over-excited for unpredicted stimuli (Fig. S2). Recently, it has been shown that the activity in the distal apical dendrites increases over time for unexpected stimuli but remains at a baseline for predictable inputs (66). While our assumptions are supported by experimental findings, we note that they can be relaxed. An excess of inhibition or excitation at the dendrites for fully predicted stimuli would not hamper the formation of PE neurons, but the learned connectivity would differ with respect to the synaptic strengths needed to establish an E/I balance at the soma.

Furthermore, we have assumed that the neurons in our network exhibit linear input-output transfer functions for positive rates. However, for real neurons, this linearity will only be true approximately for some input range. We expect that the PE circuit’s ability to withstand perturbations and to generalize to inputs not seen during learning will be compromised for neurons with strong sub- or supra-linear response functions. The network will nevertheless settle for a solution that corresponds to the linearized input-output transfer function at the range of stimuli seen during learning. Furthermore, by focusing on the steady-state responses, we have neglected the temporal characteristics of sensory stimuli. It would therefore be interesting to study the transient responses of prediction-error neurons, and their activity upon predicted and unpredicted sequential chains.

A crucial pre-requisite to learning both nPE and pPE neurons in the same network with inhibitory plasticity is to assume a homeostatic PC rate of zero. This assumption is in line with studies showing an astonishingly low spontaneous firing rate for neurons in some regions of the cortex (see, for instance, 16, 17). In fact, the existence of unidirectional PE neurons has been attributed to low baseline firing rates (1, 3). If the PCs have a baseline firing rate significantly larger than zero, nPE and pPE neurons in our network would be bi-directional. We speculate that learning nPE and pPE neurons with non-zero baseline firing rates requires different forms of plasticity rules (taking into account the type of PE neuron a PC should develop to), or gating signals that guide or restrict learning to a subset of input phases. For instance, it has been hypothesized that neuromodulators that are active during self-motion may support the formation of PE circuits (3).

In line with experimental studies (6), our network was only trained with fully predicted sensory inputs (but see Fig. S6 for examples trained with mismatch phases as well). Therefore, responses to mismatches are only implicitly constrained during learning. We found that the key factors determining whether a PC develops into an nPE neuron, pPE neuron, silent neuron or a neuron indicating mismatches independent of the PE valence are the initial connectivity before learning and the distribution of actual and predicted sensory input onto interneurons (Fig. 4). Hence, to ensure that both nPE and pPE neurons can develop, the connectivity and the distribution of inputs should be sufficiently diverse. Given that PV, SOM and VIP neurons receive a spectrum of feedforward and feedback projections (38, 43, 44), it is plausible that they receive both actual and predicted sensory inputs.

The connections onto the soma and the dendrite of PCs as well as the inhibitory connections from SOM and VIP neurons onto PV neurons underwent inhibitory plasticity. The choice of synapses that undergo experience-dependent plasticity was motivated by the observation that in layer 2/3 of V1 the excitatory neurons and PV neurons, but not SOM and VIP neurons, show experience-dependent activity (6). However, we do not expect the learning of PE neurons to be compromised when all inhibitory synapses are plastic. Recently, it has been shown that NMDA receptor-dependent plasticity in early development is crucial for the responses to unpredictable and predictable stimuli in V1 (67). This suggests that excitatory plasticity plays a pivotal role in the formation of PE neurons. While we kept all excitatory connections fixed during learning, we expect that in our network PE neurons can develop with excitatory homeostatic plasticity, inhibitory homeostatic plasticity, or both.

Here, we used two homeostatic plasticity rules, one establishing a target rate for PCs, the other a target for the total input to PCs. While both rules force the excitatory neurons to remain at baseline, only the latter one achieves an E/I balance in the PE neurons. This is a consequence of the target firing rate being equal to the rectification threshold. The objective is already satisfied when the soma is inhibited. When using a target for the total input to PCs, the plasticity rule punishes positive and negative deviations from the target equally, leading eventually to a balance of excitatory and inhibitory currents. A similar result could theoretically be achieved by simply increasing the target rate slightly. However, we speculate that learning an E/I balance in such systems would be comparatively slow as negative deviations are still bounded from below. While we have employed a homeostatic plasticity rule that establishes a target for the total input, we assume that plasticity rules processing deviations from a target membrane potential (68, 69) can be equally used to learn robust nPE and pPE neurons.

We modeled both the actual and predicted sensory stimuli as noisy step “currents” with varying strength. It would be interesting, in future work, to extend our analyses to account for the high-dimensional nature of sensory stimuli that is neglected in our simulations. In a first step, the excitatory neurons could be equipped with feature selectivity, so that they preferentially fire for a particular feature, and show only weaker responses for others. Indeed, there is evidence that neurons in layer 2/3 of rodent V1 preferentially signal the mismatch between expected and true sensory inputs in a particular location of the receptive field (70). Also, neurons in the anterior frontal-striatal cortex of macaque have been shown to encode feature-specific prediction errors (71). While excitatory neurons show pronounced feature selectivity (see, for instance, 72–75), some inhibitory neurons seem to be more broadly tuned than excitatory neurons (see, for instance, 72, 76–78). A thorough investigation of how feature selectivity in excitatory neurons influences the formation of PE neurons, and the tuning properties of different interneuron types is, however, beyond the scope of the present study.

Besides the role of prior expectations in perceptual inference (see, for instance, 1, 42, 79), prediction errors may govern learning. In the rodent visual cortex, unexpected event signals predict subsequent changes in neuron responses to predicted and unpredicted sensory stimuli (66). Recently, it has been demonstrated that there is a close link between predictive coding and supervised learning, in which non-biological weight changes by the backpropagation algorithm can be replaced with local Hebbian plasticity of connections in predictive coding networks (65, 80–82). Furthermore, it has been shown that a biologically plausible learning scheme can ease the temporal and structural credit assignment problem (83). The learning rule is based on attentional feedback signals that form synaptic tags in a subset of connections, and global neuromodulators driven by reward prediction-errors. Moreover, this model has recently received support from experimental data showing that PE neurons in anterior frontal-striatal networks could serve as feature-specific eligibility traces (71).

Expected sensory stimuli are recognized more swiftly (84, 85), giving an evolutionary advantage in a world where seconds can make the difference between life and death. Hence, continuous detection of deviations between expected and actual sensory inputs, and the subsequent refinement of predictions may be a central task for neural networks. Our work sheds light on the formation and refinement of prediction-error neurons in cortical circuits, an important step towards a better understanding of the brain’s ability to predict sensory stimuli.

## Models and methods

### Prediction-error network model

We simulated a rate-based network model of excitatory pyramidal cells (*N*_PC_ = 140) and inhibitory PV, SOM and VIP neurons (*N*_PV_ = *N*_SOM_ = *N*_VIP_ = 20). All neurons are randomly connected with connection strengths and probabilities given below (see “Connectivity”).

Each excitatory pyramidal cell is divided into two coupled compartments, representing the soma and the dendrites, respectively. The dynamics of the firing rate *r*_E,*i*_ of the somatic compartment of neuron *i* obeys (36)

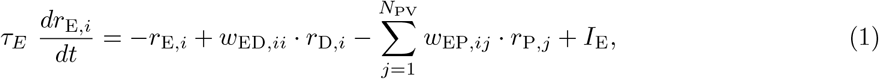

where *τ*_E_ denotes the excitatory rate time constant (*τ*_E_=60 ms), the weight *w*_ED_ describes the connection strength between the dendritic compartment and the soma of the same neuron, and *w*_EP_ denotes the strength of somatic inhibition from PV neurons. The overall input *I*_E_ comprises external background and feedforward sensory inputs (see “Inputs” below). Firing rates are rectified to ensure positivity.

The dynamics of the activity *r*_D,*i*_ of the dendritic compartment of neuron *i* obeys (36)

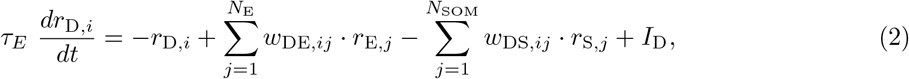

where the weight *w*_DE_ denotes the recurrent excitatory connections between PCs, including backpropagating activity from the soma to the dendrites. *w*_DS_ represents the strength of dendritic inhibition from SOM neurons. The overall input *I*_D_ comprises fixed, external background inputs and feedback predictions (see “Inputs” below). We assume that any excess of inhibition in a dendrite does not affect the soma, that is, the dendritic compartment is rectified at zero.

Just as for the excitatory neurons, the firing rate dynamics of each interneuron is modeled by a rectified, linear differential equation (36),

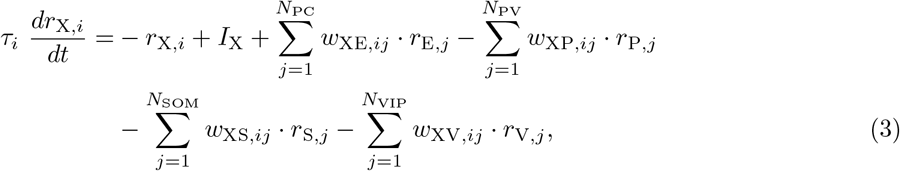

where *r*_X,*i*_ denotes the firing rate of neuron *i* from neuron type *X*, and the weight matrices *w*_XY_ denote the strength of connection between the presynaptic neuron population *Y* and the postsynaptic neuron population *X* (*X, Y* ∈ {*P, S, V*}). The rate time constant *τ*_*i*_ was chosen to resemble a fast GABA_A_ time constant, and set to 2 ms for all interneuron types included. The overall input *I*_X_ comprises fixed, external background inputs, as well as feedforward sensory inputs and feedback predictions (see “Inputs” below).

### Connectivity

All neurons are randomly connected with connection probabilities motivated by the experimental literature (e.g. 12, 13, 30–35),

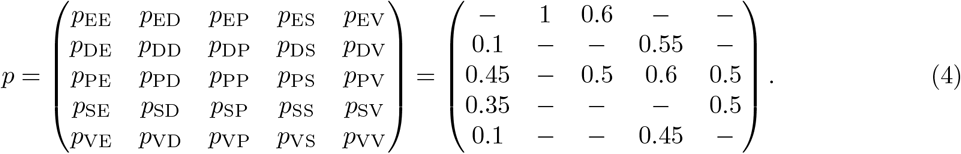

All cells of the same neuron type have the same number of incoming connections. The mean total connection strengths are set to

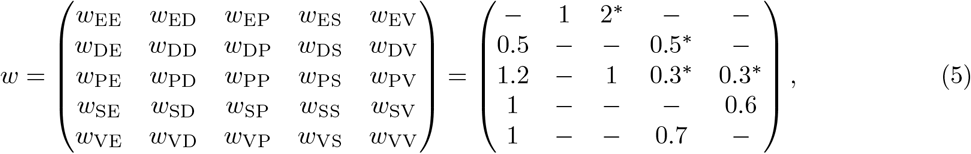

where ^*^ denotes either the weights that are free for optimization to satisfy the equations describing an E/I balance (see Supporting Information), or the initial mean connection strengths that are subject to synaptic plasticity during learning. For plastic networks, the initial connections between neurons are drawn from uniform distributions

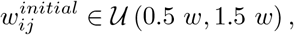

where *w* denotes the mean connection strengths given in (5). Please note that the system is robust to the choice of connection strengths. The connection strengths are merely chosen such that the solutions of the equations describing an E/I balance comply with Dale’s principle.

In plastic networks, *w*_EP_ is subdivided into assemblies. While one-third of PCs receive stronger initial inhibition from PV neurons that are driven by sensory input, another third receives stronger initial inhibition from PV neurons that are driven by feedback predictions. More precisely, for two-thirds of the excitatory neurons, half of the connections from PV neurons are strengthened by 1.5, while the remaining ones are weakened by 0.5.

All weights are scaled in proportion to the number of existing connections (i.e., the product of the number of presynaptic neurons and the connection probability), so that the results are independent of the population size.

### Inputs

All neurons receive external background input that ensures reasonable baseline firing rates in the absence of sensory inputs and predictions thereof. In the case of non-plastic networks, these inputs were set such that the baseline firing rates are *r*_E_ = 1*s*^−1^, *r*_P_ = *r*_S_ = *r*_V_ = 4*s*^−1^ and *r*_D_ = 0*s*^−1^. In the case of plastic networks, we set the external inputs of all neuron types to *x*_E_ = *x*_P_ = *x*_S_ = *x*_V_ = 5*s*^−1^, while the background input to the dendrites is dynamically computed during training such that *r*_D_ = 0*s*^−1^.

In addition to the external background inputs, the neurons receive either sensory input (*S*), a prediction thereof (*P*), or both. We distinguish between phases of fully predicted (*P* = *S >* 0), overpredicted (*P > S*) and underpredicted (*P < S*) sensory stimuli, as well as baseline phases (*P* = *S* = 0). During training, the network is exposed to baseline phases and fully predicted sensory inputs (Figs. 3 and 4), or in addition to over- and underpredicted sensory stimuli (Fig. S6). Stimuli are drawn from a uniform distribution from the interval [0, 5 *s*^−1^]. Mean and SD of test stimuli for each simulation are listed below (see “Simulations”).

### Plasticity model

In plastic networks, a number of connections between neurons are subject to experience-dependent changes in order to establish an E/I balance for PCs (see “Connectivity”). Weights were updated after the steady state firing rates have been reached, with learning rates *η*_EP_ = *η*_PS_ = *η*_PV_ = 1*e*^−3^ and *η*_DS_ = 1*e*^−4^.

### Connections onto PCs

The connections from PV and SOM neurons onto the soma and the apical dendrites, respectively, obey inhibitory Hebbian plasticity rules akin to (46)

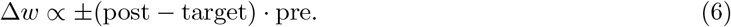

More precisely,

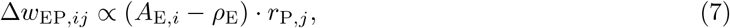

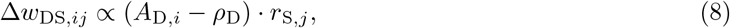

with *A*_X_ representing either the firing rate of or the total input into compartment *X* ∈ {*E, D*} (see main text). The targets *ρ*_E_ and *ρ*_D_ are both set to zero in the simulations.

### Connections onto Ins

The connections from both SOM and VIP neurons onto PV neurons implement a local approximation of a backpropagation of error rule that relies on information forwarded to the PV neurons from the PCs (23, 48).

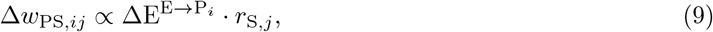

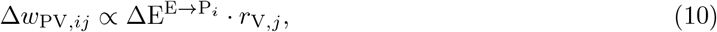

where 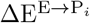 denotes the weighted difference between the excitatory recurrent drive onto the PV neuron and a target,

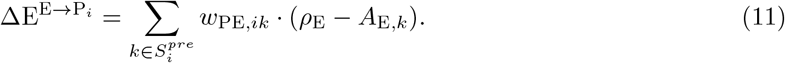

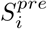 denotes the set of presynaptic PCs a particular PV neuron receives excitation from.

### Negative and positive prediction-error neurons

We define PCs as negative prediction-error (nPE) neurons when they exclusively increase their firing rate for overpredicted sensory stimuli, while remaining at their baseline for fully predicted and underpredicted sensory inputs. Similarly, we consider PCs as positive prediction-error (pPE) neurons when they exclusively increase their firing rate for underpredicted stimuli, while remaining at their baseline for fully predicted and overpredicted sensory inputs. In practice, we tolerate small deviations in phases in which the PE neurons are supposed to remain at baseline, as long as these deviations are smaller than 10% of the neuron’s maximal response. Tolerating small deviations is more in line with experimental approaches. Please note that the results do not rely on the precise thresholds used for the classification.

A detailed derivation and network analysis of the constraints imposed on the connectivity by the presence of both nPE and pPE neurons for a linearized, homogeneous network can be found in the Supporting Information. We solved eqs. (31)-(34), (36), (38), (43) and (44) to find an optimal weight configuration for a mean-field network in which SOM neurons receive the actual sensory inputs and VIP neurons receive a prediction thereof. In the same way, we solved eqs. (31)-(34), (49), (50), (53) and (55) to find an optimal weight configuration for a mean-field network in which SOM neurons receive the predicted sensory inputs and VIP neurons receive the actual sensory stimuli.

### Attractor-memory network with PE neurons

To study the effect of unpredictable events, we connected two PE circuits (see above) with two representation neurons and an attractor network generating predictions (Fig. 5). Neurons of the PE circuits receive sensory inputs randomly drawn from either of two uniform distributions. We distinguish between a distribution termed ‘weak’ ranging from 1 to 5, and a distribution termed ‘strong’ ranging from 5 to 9. All neurons of column *a* are selectively responsive to stimuli drawn from the ‘weak’ distribution, while neurons of column *b* are only responsive to stimuli drawn from the ‘strong’ distribution.

The attractor network is given by two neurons 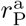 and 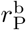 that are mutually connected via inhibitory synapses (53). Both neurons receive inputs from nPE and pPE neurons of the corresponding PE circuit, and a memory neuron, 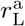 or 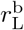,

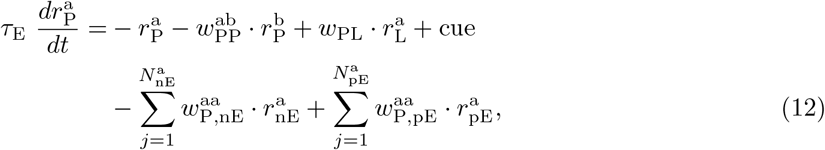

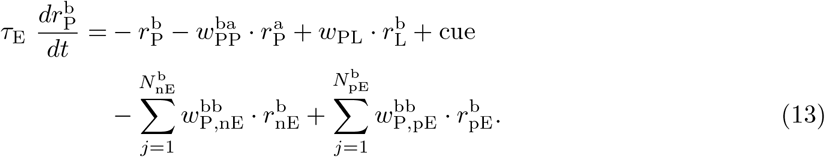

The weights *w*_PP_ denote the mutual connection strength between prediction neurons (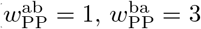, asymmetry chosen to take into account the difference between the activities of the prediction neurons), and *w w*_PL_ represents the strength of connection between the corresponding memory neuron and prediction node (*w*_PL_ = 1), and 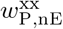 and 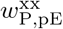 denote the strength of connection between nPE/pPE neurons and the prediction neuron, respectively 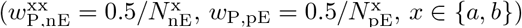. In most simulations, a cue signal indicated from which distribution the stimulus is drawn. We modeled this evidence by a strong negative input (−10*/s*) given to the prediction neuron associated with the other distribution. Firing rates are rectified to ensure positivity.

The memory neurons, 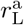 and 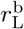, are given by two perfect integrators that obtain inputs from nPE and pPE neurons of the corresponding PE circuit,

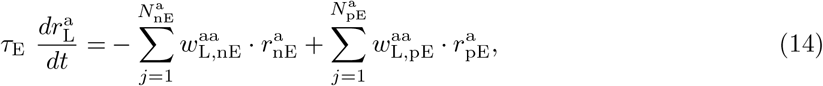

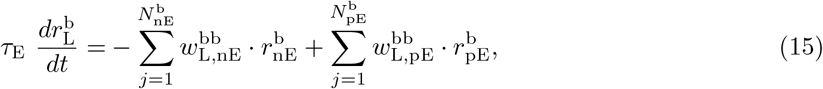

where the weights *w*_L,nE_ and *w*_L,pE_ denote the strength of connection between nPE/pPE neurons and the memory neurons 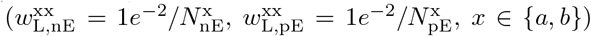. In Fig. 5 E, PE neurons of column *a* were connected to the memory neuron of column *b*, and PE neurons of column *b* were connected to the memory neuron of column *a*. This reflected the network’s knowledge about the coupling between the two distributions. In that case, we introduced the weights 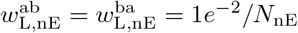 and 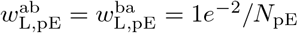 to account for the cross-coupling.

The representation neurons 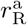 and 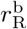 receive the sensory stimulus *S* and connections from PE neurons of that column,

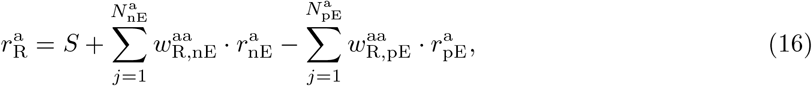

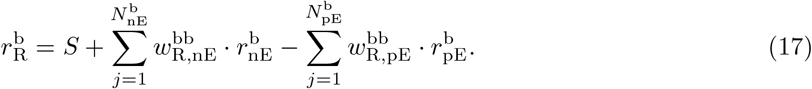

The weights *w*_R,nE_ and *w*_R,pE_ represent the strength of connections between the nPE/pPE neurons and the output neuron, respectively 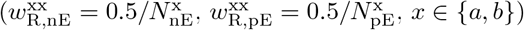.

### Simulations

All simulations were performed in customized Python code written by LH. Differential equations were numerically integrated using a 2^nd^-order Runge-Kutta method with time steps ranging between 0.1 and 0.2 ms. Neurons in the PE circuits were initialized with *r* = 0*/s*. The memory neurons were initialized at the mean of the two distributions (see above), and each prediction neuron was either set to the mean of the distribution it is associated with if the stimulus at *t* = 0 ms was drawn from that distribution, or set to zero otherwise. Each stimulus was presented for 1 second. During training of the PE circuit, we presented 350 stimuli alternated with 350 zero-input (baseline) phases. We made sure that the weights converged to a configuration that satisfied the objective given by our homeostatic plasticity rules (see Eqs. 7-10 in “Plasticity model”). We defined the steady-state firing rate per stimulus as the activity in the last 500 ms of stimulus presentation. The onset firing rate was computed as the activity of the first 10 ms.

**Figures 1 & S2:** Test stimulus was set to 5*/s* with a SD of 1*/s*. Stimulus to compute total excitatory and inhibitory inputs was set to 1*/s*.

**Figures 2 & S3:** Test stimulus was set to 3*/s* with a SD of 1*/s*. The perturbation stimuli ranged from −1.5*/s* to 1.5*/s*.

**Figures 3, S4 & S5:** Test stimulus was set to 5*/s* with a SD of 1.5*/s*. 50% of the PV neurons, 70% of the SOM neurons and 30% of the VIP neurons receive the actual sensory input, while the remaining ones of each population received a prediction thereof. Perturbation stimulus was *±*2*/s*. Panels D & G of Fig. 3 show the median over all PE neurons and the SEM.

**Figures 4 & S6:** Test stimulus was set to 5*/s* with a SD of 1.5*/s*. In main figure, square: *w*_EP_ ∈ [2, 4], *w*_PS_ ∈ [0.5, 1], *w*_PV_ ∈ [1.5, 2.5]; circle: *w*_EP_ ∈ [2.5, 8], *w*_PS_ ∈ [1.5, 2.5], *w*_PV_ ∈ [0.5, 1]; triangle: *w*_EP_ ∈ [2.5, 8], *w*_PS_ ∈ [1, 2.5], *w*_PV_ ∈ [0.5, 2]. Half of the PV neurons and all SOM neurons receive the actual sensory input, while the remaining PV and SOM neurons as well as all VIP neurons receive a prediction thereof. In supporting figure: Total number of stimuli presented during training was increased, so that the number of fully predicted sensory stimuli was constant at 350. Results were averaged over 5 simulations, mean and SD are shown.

**Figure 5:** For panel E, the performance error was computed as the squared difference between the activity of the respective line attractor and the stimulus presented. For panel F, the initial weight between the stimulus and the representation neuron was set to 0.5. The basis learning rate (fixed) was set to 5*e*^−4^. And the initial speed was computed as the derivative of the rate with respect to time, averaged over the first 50 ms.

Source code to reproduce the simulations, analyses and figures will be available after publication at github.com/lhertaeg/SourceCode_Hertaeg2021.

## Acknowledgments

We are grateful to Vezha Boboeva, Douglas Feitosa Tomé, Júlia Gallinaro and Klara Kaleb for helpful comments on earlier versions of this manuscript, and we want to thank all members of the Clopath lab for discussion and support. This work was supported by BBSRC BB/N013956/1, BB/N019008/1, Wellcome Trust 200790/Z/16/Z, Simons Foundation 564408 and EPSRC EP/R035806/1.

## Supplementary Information

### Deriving constraints for the interneuron circuit imposed by the presence of both nPE and pPE neurons

We performed a mathematical analysis of a simplified model to identify the constraints that are imposed on the interneuron circuit by the simultaneous presence of both nPE and pPE neurons. In this analysis, we assume that all neuron types exhibit a sufficiently high baseline firing rate so that the rectifying nonlinearities for PCs, PV, SOM, and VIP neurons can be neglected. Merely the rectification in the dendrites is retained so that any excess of inhibition in a dendrite does not affect the soma. Hence, we will perform a case differentiation in the subsequent analysis. These assumptions reduce the network to a system of analytically tractable linear equations.

We furthermore consider a homogeneous network, that is, all weights, neuronal properties and the number of incoming connections for cells of the same type are the same. As a result, we can reduce the high-dimensional system to a low-dimensional set of linear equations, each equation describing the dynamics of one representative firing rate per neuron type/compartment,

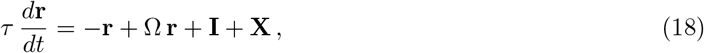

where *τ* denotes the rate time constant, the vector **r** represents the activity level for each neuron type/compartment, Ω contains the recurrent strength of connections between neurons/compartments, and **X** and **I** denote the external background input and the actual/predicted sensory inputs, respectively. In the steady state, the firing rates are given by

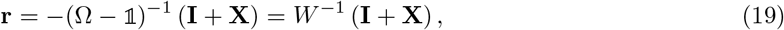

with the effective connectivity matrix *W* that includes the leak,

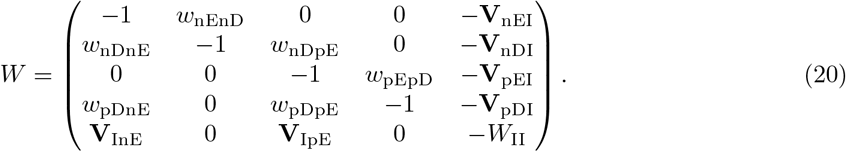

**V**_InE_ and **V**_IpE_ are column vectors of length *N*_I_. They contain the effective weights from nPE and pPE neurons to all interneurons I, respectively. **V**_nEI_, **V**_nDI_, **V**_pEI_ and **V**_pDI_ are row vectors of length *N*_I_, and comprise the connection strengths from all interneurons I to the soma and dendrites of nPE and pPE neurons. Furthermore, the squared matrix *W*_II_ denotes the effective connectivity between all interneuron types. All weights between neuron types are strictly positive to maintain the excitatory/inhibitory nature of the various cell classes considered here.

The input **I** contains the sensory feedforward input *S* and the prediction *P* thereof for each neuron type/compartment *Y*, 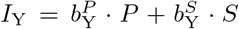 (the parameters 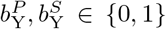 control different input configurations and characterize the phases). The baseline phase is characterized by the absence of feedforward sensory inputs *S* and feedback predictions *P* (**I** = **0**). In contrast, for fully predicted stimuli, sensory inputs and predictions are the same (**I** = *s ·* 𝟙_FP_, with 𝟙_FP_ a vector of 1’s and *s* ≥ 0). For underpredicted stimuli, the sensory input is larger than the prediction thereof. For the sake of convenience, we set *P* to zero, hence, **I** = *s ·* 𝟙_UP_ (with 𝟙_UP_ a vector of zeros and ones, and *s* ≥ 0). Last but not least, for overpredicted stimuli, the sensory input is smaller than the prediction thereof. As before, for the sake of convenience, we set *S* to zero, hence, **I** = *s ·* 𝟙_OP_ (with 𝟙_OP_ a vector of zeros and ones, and *s* ≥ 0).

To be considered as a nPE neurons, the soma of PCs must remain at baseline for fully predicted and underpredicted stimuli, while increasing activity relative to baseline for overpredicted stimuli,

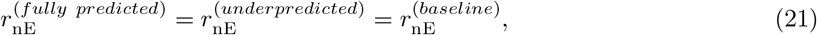

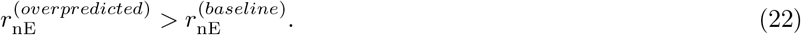

Similarly, an excitatory neuron is classified as a pPE neuron when its activity remains at baseline for fully predicted and overpredicted sensory stimuli, while increasing activity relative to baseline for underpredicted stimuli,

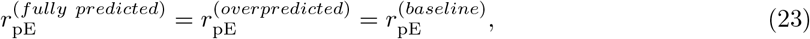

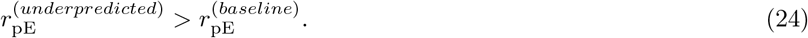

In the steady state, eqs. (19) and (20) yield

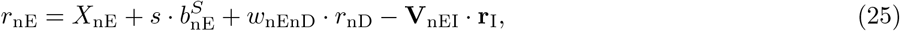

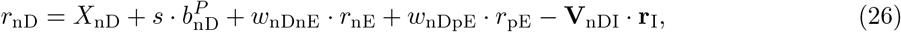

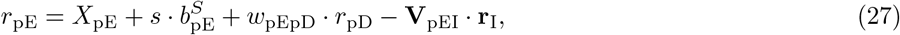

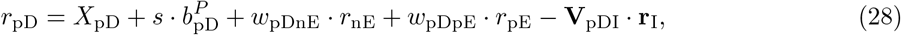

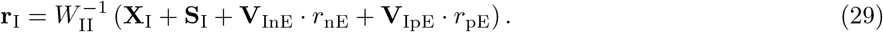

While nPE and pPE neurons remain at their baseline for perfectly predicted stimuli (eqs. 21 and 23), the inhibitory neurons change their activity,

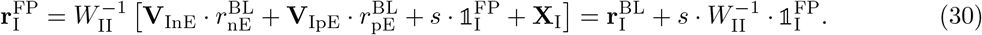

We assume that the dendrites of nPE and pPE neurons also remain at their baseline for fully predicted stimuli, set to zero for mathematical convenience. Together with eq. (30), eqs. (25)-(28), hence, yield

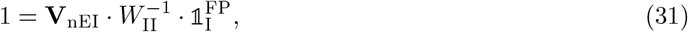

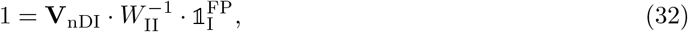

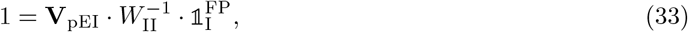

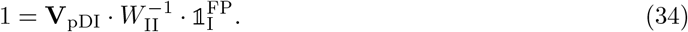

#### Case I: Dendrites are inhibited for underpredicted and excited for overpredicted stimuli

##### Underprediction

While nPE neurons remain at baseline for sensory inputs that are larger than predicted (eq. 21), pPE neurons increase their activity relative to baseline (eq. 24)

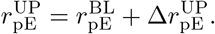

The steady state firing rate of interneurons for underpredicted stimuli is then given by

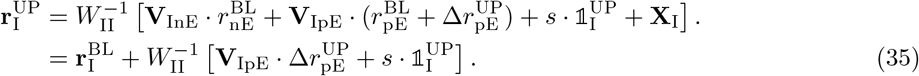

Inserting this eq. into eq. (27) yields

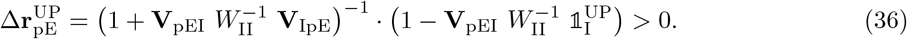

Inserting, eq. (35) into eq. (25) yields

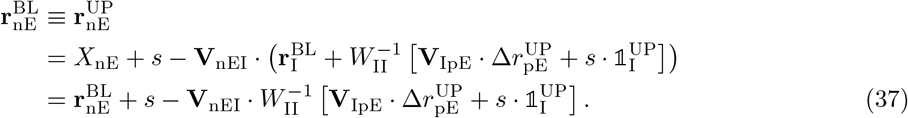

Together with eq. (36), the condition reads

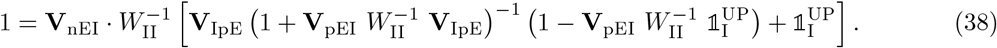

##### Overprediction

While pPE neurons remain at baseline for sensory inputs that are smaller than predicted (eq. 23), nPE neurons increase their activity relative to baseline (eq. 22)

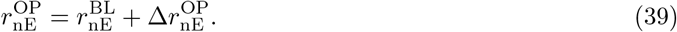

The steady state firing rate of interneurons during overprediction is then given by

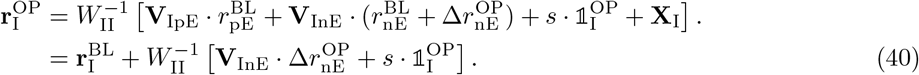

While dendrites could be neglected for underpredicted stimuli, they must be considered for overpredicted stimuli. Inserting eqs. (39) and (40) into eqs. (26) and (28) yields

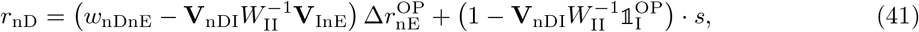

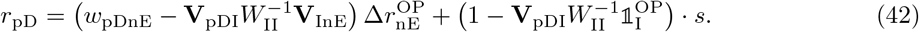

Inserting eqs. (40) and (41) into eq. (25) gives

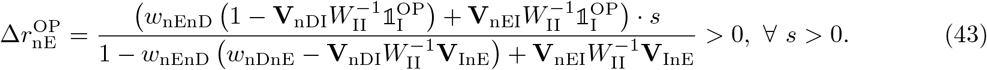

pPE neurons remain at baseline for overpredicted stimuli. By inserting eqs. (40) and (42) into eq. (27), we obtain

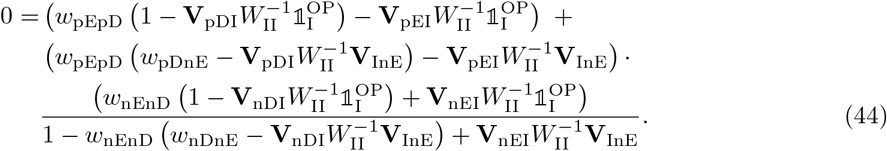

#### Case II: Dendrites are excited for underpredicted and inhibited for overpredicted stimuli

##### Underprediction

While nPE neurons remain at baseline for sensory inputs that are larger than predicted (eq. 21), pPE neurons increase their activity relative to baseline (eq. 24)

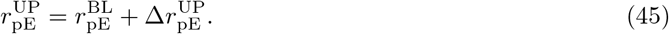

The steady state firing rate of interneurons for underpredicted stimuli is then given by

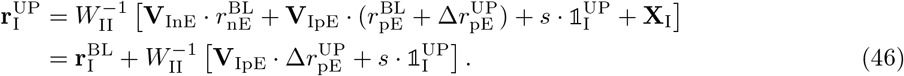

While the dendrites can be neglected for overpredicted stimuli, they must be considered for underpredicted stimuli. Inserting eqs. (45) and (46) into eqs. (26) and (28) yields

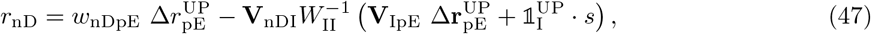

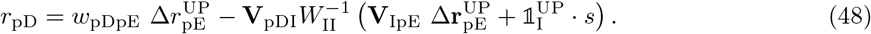

Inserting eqs. (46) and (48) into (27) gives

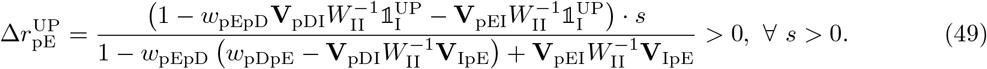

nPE neurons remain at baseline for underpredicted sensory inputs. By inserting eq. (46) and (47) into eq. (25), we obtain

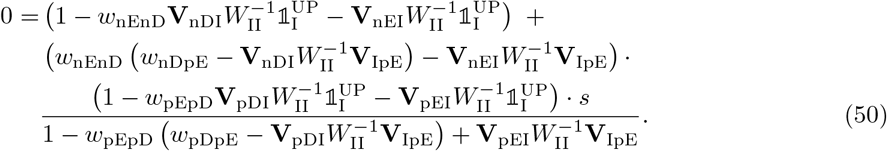

##### Overprediction

While pPE neurons remain at baseline for sensory inputs that are smaller than predicted (eq. 23), nPE neurons increase their activity relative to baseline (eq. 22)

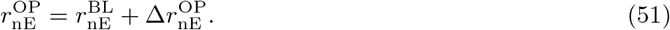

The steady state firing rate of interneurons for overpredicted stimuli is then given by

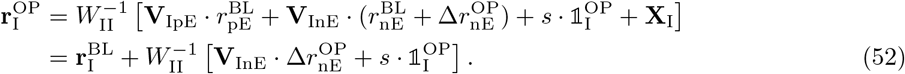

Inserting this into eq. (25) yields

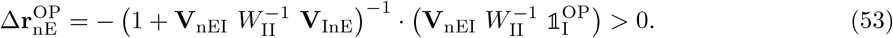

Inserting eq. (52) into eq. (27) yields

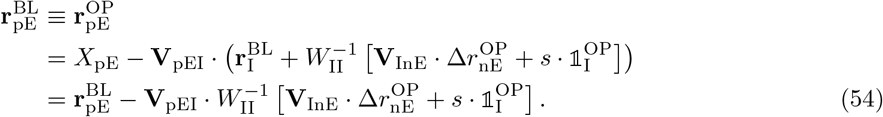

Together with eq. (53), the condition reads

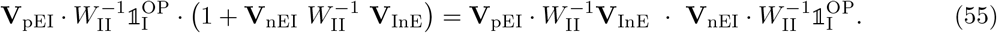

### The presence of both nPE and pPE neurons requires diversified somatic inhibition

We consider an interneuron circuit with one PV, SOM and VIP population. As before, we assume that we can represent the steady state activity of each population by one representative per neuron type (that is, a mean-field approach). While the PV neuron provides somatic inhibition, the SOM neuron provides dendritic inhibition (11, 14). Consequently, **V**_nEI_, **V**_nDI_, **V**_npI_ and **V**_pDI_ read

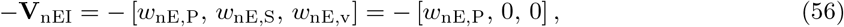

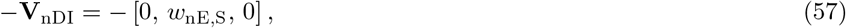

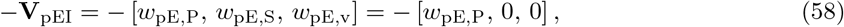

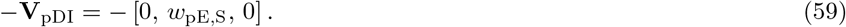

In the following, we consider the case in which dendrites are inhibited for underpredicted stimuli and excited for overpredicted stimuli. For underpredicted inputs, eq. (36) gives

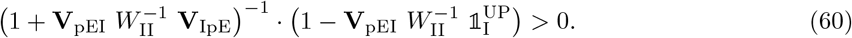

Without loss of generality, we assume that 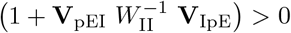. Hence, the preceding equation simplifies to

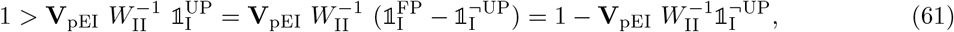

where 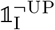 denotes a vector that contains zeros for each neuron that receives sensory input, ones otherwise. Hence, we get

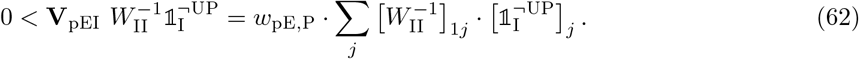

As *w*_pE,P_ *>* 0, we deduce that

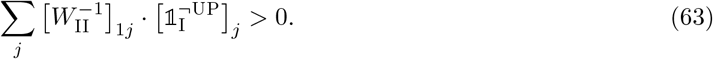

The condition for nPE neurons for underpredicted stimuli, eq. (38), can be written as

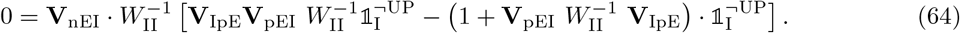

Because some of the terms cancel, the equation simplifies to

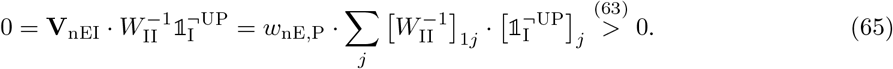

This contradiction shows that the somatic inhibition provided by PV neurons must be diversified in order to satisfy the constraints imposed by the simultaneous presence of nPE and pPE neurons. The same logic applies to the case in which dendrites are excited for underpredicted inputs and inhibited for overpredicted stimuli.

### PE circuits with one PE neuron type

PE circuits with either nPE or pPE neurons require only one source of somatic inhibition. We chose the interneuron connectivity according to a well-known motif that has been observed in many brain areas, including primary visual cortex (12, 13, 32),

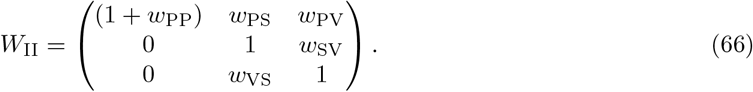

The inverse is directly given by

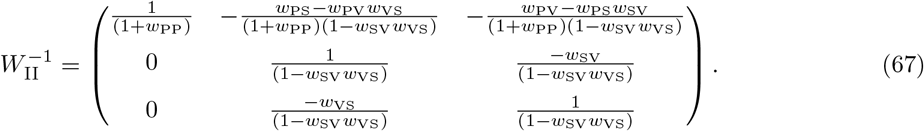

#### PE circuits with nPE neurons only

With the interneuron connectivity described above and eqs. (31) - (32), excitatory neurons remain at their baseline for perfectly predicted stimuli when

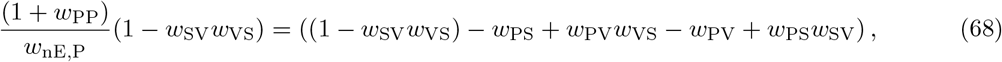

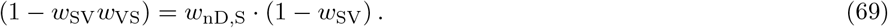

If SOM neurons receive sensory inputs, and VIP neurons receive either actual or predicted sensory inputs, the dendrites are inhibited for underpredicted inputs and excited for overpredicted stimuli. As pPE neurons are not present, eq. (38) simplifies to

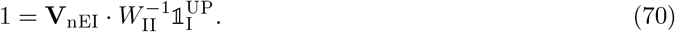

If SOM neurons receive the predicted sensory inputs, and VIP neurons receive the actual sensory inputs, the dendrites are excited for underpredicted sensory stimuli and inhibited for overpredicted inputs. As pPE neurons are not present, eq. (50) simplifies to

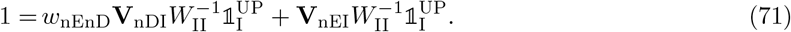

#### PE circuits with pPE neurons only

With the interneuron connectivity described above and eqs. (33) - (34), excitatory neurons remain at their baseline for perfectly predicted stimuli when

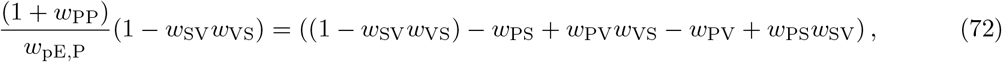

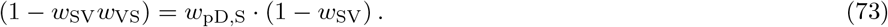

If SOM neurons receive sensory inputs, and VIP neurons receive either actual or predicted sensory inputs, the dendrites are inhibited for underpredicted stimuli and excited for overpredicted inputs. As nPE neurons are not present, eq. (44) simplifies to

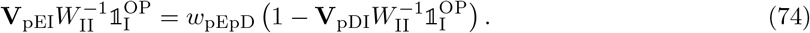

If SOM neurons receive the predicted sensory inputs, and VIP neurons receive the actual sensory inputs, the dendrites are excited for underpredicted sensory inputs and inhibited for overpredicted inputs. As nPE neurons are not present, eq. (55) simplifies to

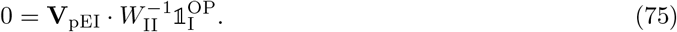

## Supplementary Figures

**Figure S1.**
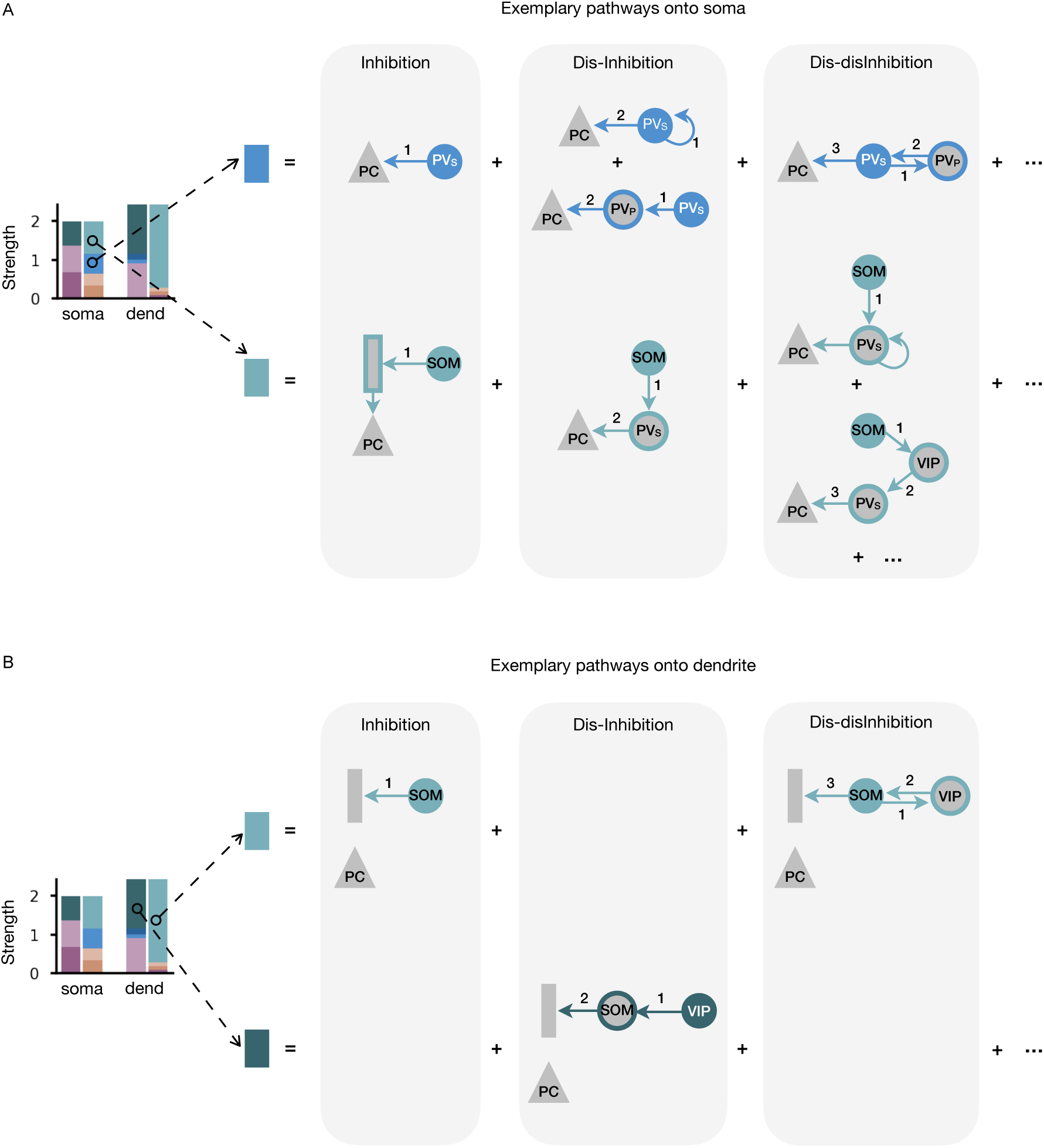
Illustration of pathways onto soma and dendrites of PE neurons. **(A)** For each neuron type/compartment, the strengths of all pathways that originate from this node and end at the soma of a PE neuron are summed up. The example pathways depicted comprise a number of inhibitory, dis-inhibitory and dis-disinhibitory pathways. Numbers 1 to 3 denote input flow through the circuit after arriving at the particular neuron type/compartment (filled, color-coded). Ellipses indicate pathways that are not shown for clarity. **(B)** Same as in A but for pathways ending at the dendrites of PE neurons.

**Figure S2.**
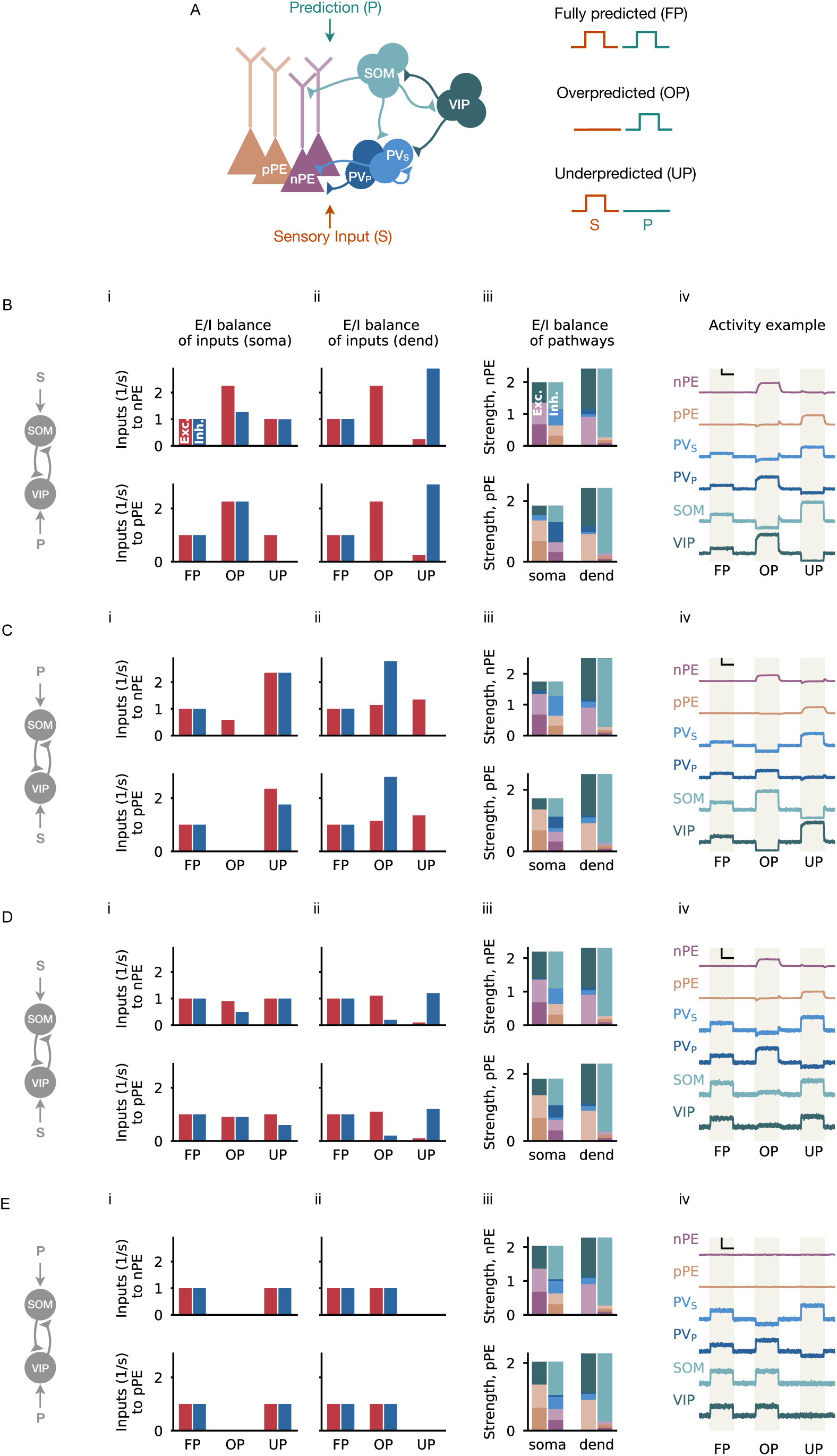
Multi-pathway E/I balance in nPE and pPE neurons for different PE circuits. **(A)** Illustration of a network model with excitatory PCs and inhibitory PV, SOM and VIP neurons. Connections from PCs not shown for the sake of clarity. In a PE circuit, PCs either act as nPE or pPE neurons. Somatic compartment of PCs and half of the PV neurons receive the actual sensory input, while the dendritic compartment of PCs and the remaining PV neurons receive the predicted sensory input. The inputs to SOM and VIP neurons are varied (B-E). Sensory stimuli can either be fully predicted (FP), overpredicted (OP) or underpredicted (UP). **(B)** In a mean-field network in which SOM neurons receive the actual sensory input and VIP neurons receive a prediction thereof, the excitatory (red) and inhibitory (blue) inputs at the soma (i) and at the dendrite (ii) are balanced for fully predicted sensory stimuli for both nPE (top) and pPE neurons (bottom). At the soma, this balance is preserved for underpredicted stimuli (nPE neurons) or overpredicted stimuli (pPE neurons). Stimulus strength is 1 *s*^−1^. (iii) Both nPE and pPE neurons exhibit balanced pathways onto soma and dendrites. For each neuron type/compartment, the net pathway strength is the sum of all pathways that originate from this node and end either at the soma or the dendrites of nPE and pPE neurons, respectively. Color-code from A. (iv) Example network parameterized to establish an E/I balance in the pathways. Vertical black bar denotes 3 *s*^−1^, horizontal black bar denotes 500 ms. **(C)** Same as in A but for a mean-field network in which SOM neurons receive the predicted sensory input while VIP neurons receive the actual sensory input. **(D)** Same as in A but for a mean-field network in which both SOM and VIP neurons receive the actual sensory input. **(E)** Same as in A but for a mean-field network in which both SOM and VIP neurons receive the predicted sensory input.

**Figure S3.**
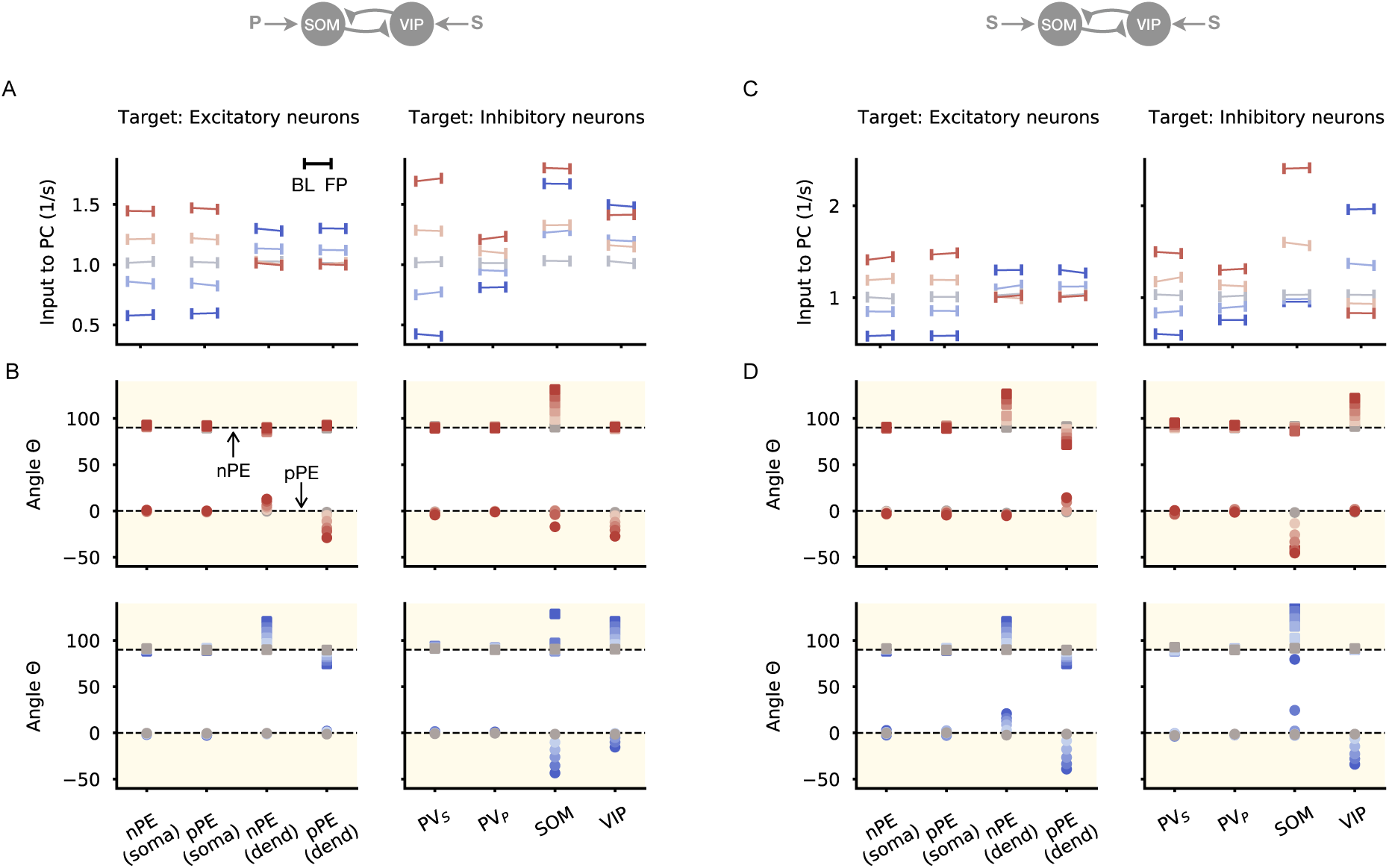
PE circuits with different inputs onto SOM and VIP neurons are robust to moderate network perturbations. **(A-D)** Each neuron type/compartment of a mean-field network with nPE and pPE neurons are perturbed with an additional inhibitory or excitatory input. Same circuit as in Fig. 1 but inputs onto both SOM and VIP neuron are varied. (A-B): SOM neurons receive the predicted sensory stimulus and VIP neurons receive the actual sensory stimulus. (C-D): Both SOM and VIP neurons receive the actual sensory input. **(A)** Total input into PE neurons during the absence of sensory stimuli (baseline, BL) and for fully predicted sensory stimuli (FP) for different perturbation strengths and different perturbation targets (Left: compartments of PCs, Right: inhibitory neurons). The total inputs in both phases are almost equal. **(B)** Θ for different perturbation strengths (top: excitatory, bottom: inhibitory) and different perturbation targets (Left: compartments of PCs, Right: inhibitory neurons). Perturbations have minor effects on nPE and pPE neurons, especially when baseline firing rates of PCs are low. **(C)** Same as in A but for a network in which both SOM and VIP neurons receive the actual sensory input. **(D)** Same as in B but for a network in which both SOM and VIP neurons receive the actual sensory input.

**Figure S4.**
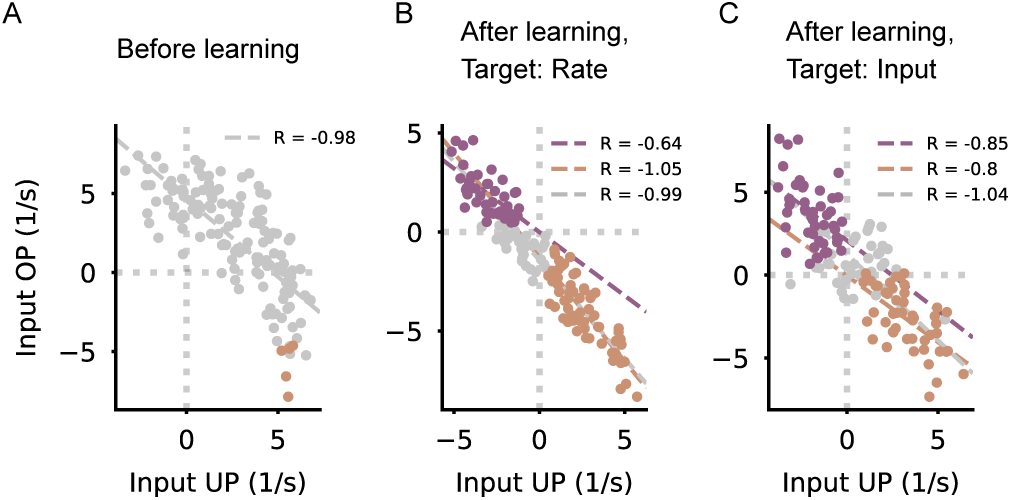
Balance of total PC inputs for actual and predicted sensory stimuli before and after learning. **(A)** Total input into PCs for overpredicted (OP) and underpredicted (UP) stimuli before learning. Individual dots denote individual neurons. **(B)** Same as in A but after learning with an inhibitory plasticity rule that establishes a zero target rate in PCs. Balance is preserved. **(C)** Same as in A but after learning with an inhibitory plasticity rule that establishes a target for the total input in PCs. Balance is preserved.

**Figure S5.**
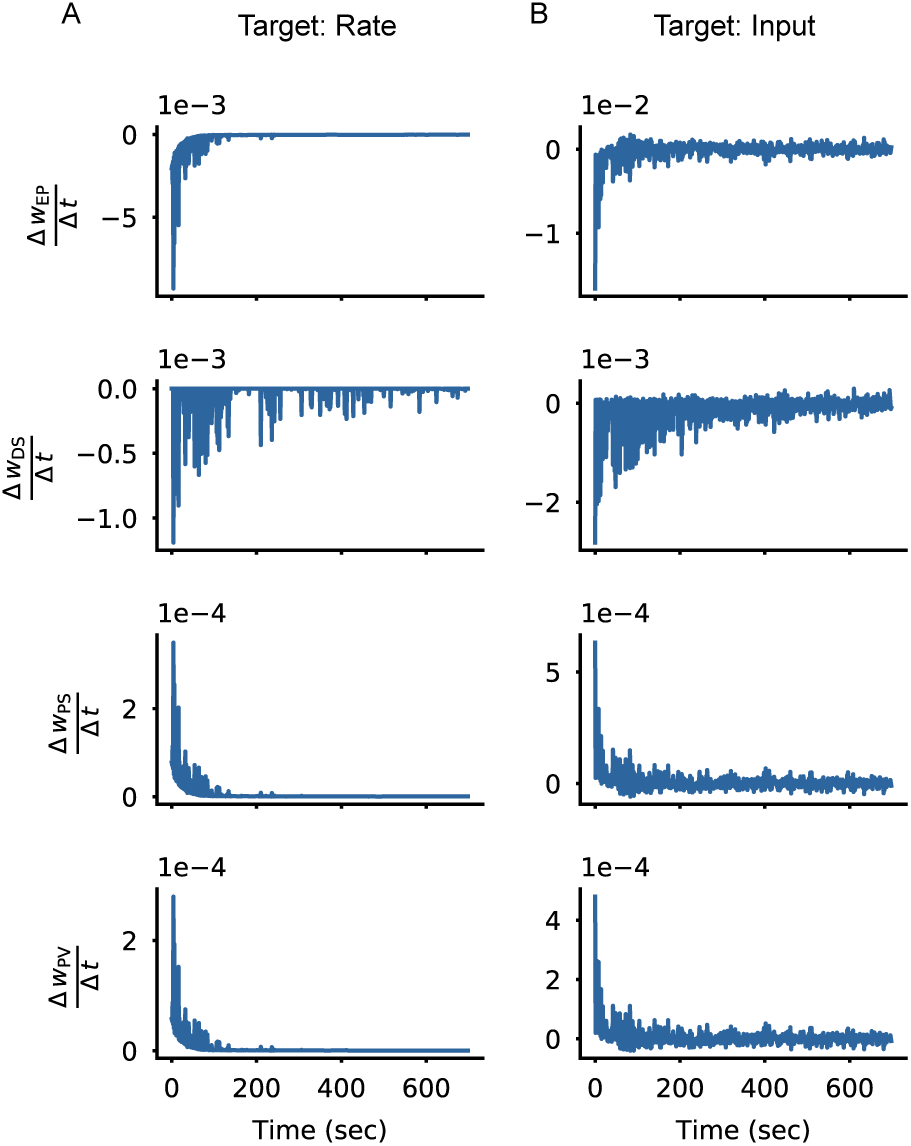
Convergence of plastic weights. **(A)** The derivatives of weights with respect to time converge to zero for an inhibitory plasticity rule that establishes a zero target rate in PCs (top to bottom: weight from PV neurons onto the soma of PCs, weight from SOM neurons onto the dendrites of PCs, weight from SOM neurons onto PV neurons, and weight from VIP neurons onto PV neurons). **(B)** Same as in A but for an inhibitory plasticity rule that establishes a target for the total input in PCs (target: zero).

**Figure S6.**
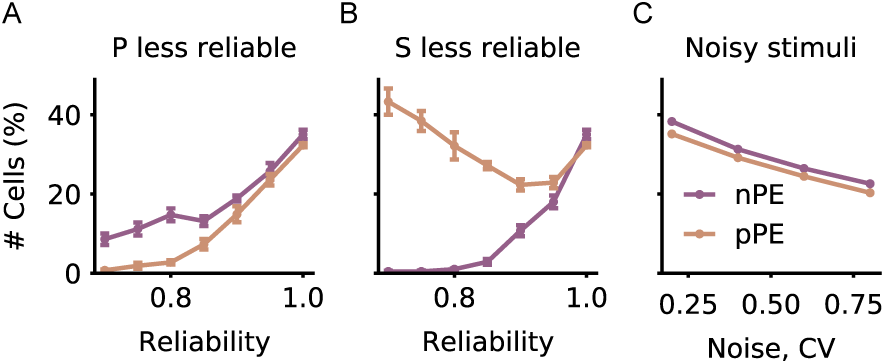
Predictability of stimuli during learning biases the formation of nPE and pPE neurons. **(A)** Number of nPE (purple) and pPE neurons (orange) decreases with decreasing reliability of the predicted sensory input. Number of pPE neurons decreases faster leading to a biased ratio of nPE and pPE neurons in favor of nPE neurons. **(B)** Number of nPE neurons decreases while the number of pPE neurons increases with decreasing reliability of the actual sensory input, leading to a biased ratio of nPE and pPE neurons in favor of pPE neurons. **(C)** Number of both nPE and pPE neurons decreases with increasingly noisy stimuli but remains at a high level for a wide range of noise levels.

